# The Influence of Angiotensin II on the Gut Microbiome of Mice: Findings from a Retrospective Study

**DOI:** 10.1101/2023.04.08.536089

**Authors:** Rikeish R. Muralitharan, Michael E. Nakai, Matthew Snelson, Tenghao Zheng, Evany Dinakis, Liang Xie, Hamdi Jama, Madeleine Paterson, Waled Shihata, Flavia Wassef, Antony Vinh, Grant R. Drummond, David M. Kaye, Charles R. Mackay, Francine Z. Marques

**Author notes:** **Correspondence to:** A/Prof Francine Marques. Hypertension Research Laboratory, School of Biological Sciences, Monash University, 25 Rainforest Walk, Clayton, Victoria 3800, Australia. E.

## Abstract

**Introduction:** Animal models are regularly used to test the role of the gut microbiome in hypertension. Small-scale pre-clinical studies have investigated changes to the gut microbiome in the angiotensin II hypertensive model. However, the gut microbiome is influenced by internal and external factors not regularly considered in the study design. Once these factors are accounted for, it is unclear if microbiome signatures are reproduceable. We aimed to determine the influence of angiotensin II treatment on the gut microbiome using a large and diverse cohort of mice and to quantify the magnitude by which other factors contribute to microbiome variations.

**Methods and Results:** We conducted a retrospective study to establish a diverse mouse cohort resembling large human studies. We sequenced the V4 region of the 16S rRNA gene from 538 samples across the gastrointestinal tract of 303 male and female C57BL/6J mice randomised into sham or angiotensin II treatment from different genotypes, diets, animal facilities, and age groups. Analysing over 17 million sequencing reads, we observed that angiotensin II treatment influenced α-diversity (P=0.0137) and β-diversity (i.e., composition of the microbiome, P<0.001). Bacterial abundance analysis revealed patterns consistent with a reduction in short-chain fatty acid-producers, microbial metabolites that lower blood pressure. Furthermore, animal facility, genotype, diet, age, sex, intestinal sampling site, and sequencing batch had significant effects on both α- and β-diversity (all P<0.001). Sampling site (6.8%) and diet (6%) had the largest impact on the microbiome, while angiotensin II and sex had the smallest effect (each 0.4%).

**Conclusions:** Our large-scale data confirmed findings from small-scale studies that angiotensin II impacted the gut microbiome. However, this effect was modest relative to most of the other factors studied. Accounting for these factors in future pre-clinical hypertensive studies will increase the likelihood that microbiome findings are replicable and translatable.

## Introduction

In the last decade, the gut microbiome has emerged as a novel factor contributing to many health and pathological states, including hypertension. Small scale case-control microbiome studies are regularly conducted in animal models – these are usually used as a proof-of-concept to show association between the gut microbiome and the disease of interest. However, regularly these studies fail to account for internal, external and technical factors that often co-influence the gut microbiome such as diet, living environment including breeding facilities, and cohort effects (Table S1).^1–11^ These factors, if not accounted or adjusted for, could result in non-biologically relevant findings that are not replicable nor translatable. We know these factors influence the gut microbiome to a certain extent from previous smaller studies,^3^ but the combined magnitude of their impact is unclear. Similarly, human gut microbiome studies are often affected by many factors which are nearly impossible to control at an experimental level. These external factors are often adjusted using normalization methods during data analysis.^12^ However, small-scale animal studies are usually under-powered to perform similar analyses.

The angiotensin II model is the most commonly used pre-clinical hypertensive model.^13^ In small-scale studies, angiotensin II treatment has previously been shown to influence the gut microbiome in mice.^14–16^ Here, we aimed to replicate the design of human populational studies in mice by performing a large retrospective study to determine the impact of angiotensin II on the gut microbiome. In our retrospective study, we included mouse samples from different diets and animal house facilities, ages, sex, cohorts, genotypes, varied doses of angiotensin II (copying varied blood pressure levels as in humans). Using this heterogenous study cohort, we were able to determine what are the most important factors that influence the microbiome in an experimental hypertensive setting and provide robust evidence for an association between angiotensin II and specific microbial taxa.

## Methods

### Animals

Male and female mice [wild-type C57BL/6J (genotype 1), and knockout genotypes developed on the C57BL/6J background GPR41/43/109aKO (genotype 2), GPR41/43KO (genotype 3), and GPR65KO (genotype 4)], from three animal house facilities in Melbourne, Australia (Animal Facility 1, Animal Facility 2, and Animal Facility 3), were included in this study (n=302; 253 male and 49 female mice). This is a retrospective study and samples used came from independent studies where animal ethics approval was previously received. Mice were administered different diets [control 1 (AIN93G, Speciality Feeds); control 2 (chow, Barastoc); high-fibre (SF11-025, Speciality Feeds); no-fibre (SF09-028, Speciality Feeds); and high-amylose corn starch acetylated and butyrylated (30% HAMSAB, Ingredion, added to AIN93G)]. The only difference between the control 1, high-fibre, and no-fibre diets is the percentage of fibre, while protein, carbohydrate and fat content are similar across the diets. Some littermate mice were randomised and underwent minipump surgery containing angiotensin II (0.5 or 0.75 mg/kg body weight/day, n=142) or 0.9% sodium chloride (sham, n=45). Gut microbiota samples were collected from young (10-12 weeks, n=264) and aged (6 months, n=38) mice. Gut bacterial samples were collected from the caecum for all samples. Gastrointestinal contents for sequencing were also collected from the small intestine (separated into duodenum, jejunum, ileum) and large intestine (caecum, colon, rectum, faeces) from some mice (n=13). The gastrointestinal contents were snap-frozen in liquid nitrogen and stored at −80°C until further processing.

### DNA Extraction, Library Preparation, and Sequencing

All samples were processed in the same laboratory. DNA was extracted using the DNeasy PowerSoil Kit (Qiagen) according to the manufacturer’s protocol. The V4 region of the bacterial 16S rRNA was amplified by PCR (Veriti Thermal Cycler, ThermoFisher Scientific) using 20 ng of DNA, Platinum Hot Start PCR master mix (ThermoFisher Scientific), 515F and 806R primers (Bioneer), single-indexing and methods as described in the Earth Microbiome Project.^17^ Thus, the microbiome data reported in this study refers to amplicon sequencing data. The quantity and quality of the PCR product were determined using a Qubit (ThermoFisher Scientific). To identify contamination, non-template controls were used, and none showed amplification. PCR products of 240ng per sample were pooled and cleaned using the PureLink PCR Purification kit (ThermoFisher Scientific). Sequencing was performed across six batches (Mar-20, Oct-20, May-21, Aug-21, Dec-21, Nov-22) in an Illumina MiSeq sequencer, generating 300 bp paired-end reads with 20% PhiX spiked in.

### Bioinformatic and Statistical Analyses of Gut Microbiome

We used the QIIME2 framework to analyse the sequence reads.^18^ The forward and reverse reads were truncated at base number 220 for forward and 200 for reverse reads. The reads were then denoised, merged, and filtered for chimera using the DADA2 plugin, resulting in an amplicon sequence variant (ASV) table at the single-nucleotide level.^19^ ASVs were then labelled by taxonomic assignment using a naïve Bayes classifier (via q2-feature-classifier),^20^ using the SILVA database (version 138) as a reference.^21^ The resulting taxonomic table was then uploaded to MicrobiomeAnalyst for downstream analyses.^22,23^ The data table was first filtered for low prevalence (<20%) and low variance (10% of interquartile range) reads. Data were then normalised using trimmed mean of M-values (TMM). The normalized data table was then used to generate two important indices commonly described in gut microbiome studies, the α- (a within-sample index) and β-diversity (a between-sample index). The α-diversity measure was determined using Shannon index, which is a more comprehensive metrics that determines both evenness and richness. α-diversity is better explained by taking into consideration both evenness and richness compared to metrics which only consider evenness or richness; thus, Shannon index was chosen. Shannon index data were regraphed in GraphPad Prism version 9 and outliers removed using robust regression and outlier removal (1%). Two-tailed Mann-Whitney U test (for 2 groups) or Kruskal-Wallis (for 3 or more groups), with adjustment for false discovery rate (FDR) was performed. SPSS for Windows (release 25) was used for sensitivity analyses to determine which factors impacted Shannon index. A step-wise multiple linear regression model was used with all the factors as independent parameters (criteria of F-entry probability: 0.05, removal: 0.10). β-diversity was determined using Bray-Curtis index, which takes into account the composition and abundance of the taxa. Differences between groups were assessed using permutational multivariate analysis of variance (PERMANOVA) for the overall effect. P-values < 0.05 were considered significant.

MaAsLin2 (in-built into MicrobiomeAnalyst) was used to determine multivariable association between taxa counts and the various experimental factors by generalized linear regression model using TMM normalized data.^24^ For the discovery of differentially abundant taxa, adjustment was made for animal house facility, genotype, diet, age, sex, sample acquisition site, treatment with angiotensin II (hypertension phenotype), and sequencing batch. FDR-adjusted P-values of <0.05 were considered significant.

To determine the explained variance of each experimental factor on the gut microbiota composition in our study, we used VpThemAll.^25^ This package enabled us to determine the full, shared, and unique variance contributed by the eight experimental factors in our study. FDR-adjusted P-values of <0.05 were considered significant. Adjusted r-squared values that were obtained were converted into percentage of variance.

## Results

### Sequencing reads and depth

Over 17,809,928 reads were denoised, merged, and survived chimera filtering, with read counts averaging 31,410 reads per sample (Figure 1a,b).

**Figure 1.**
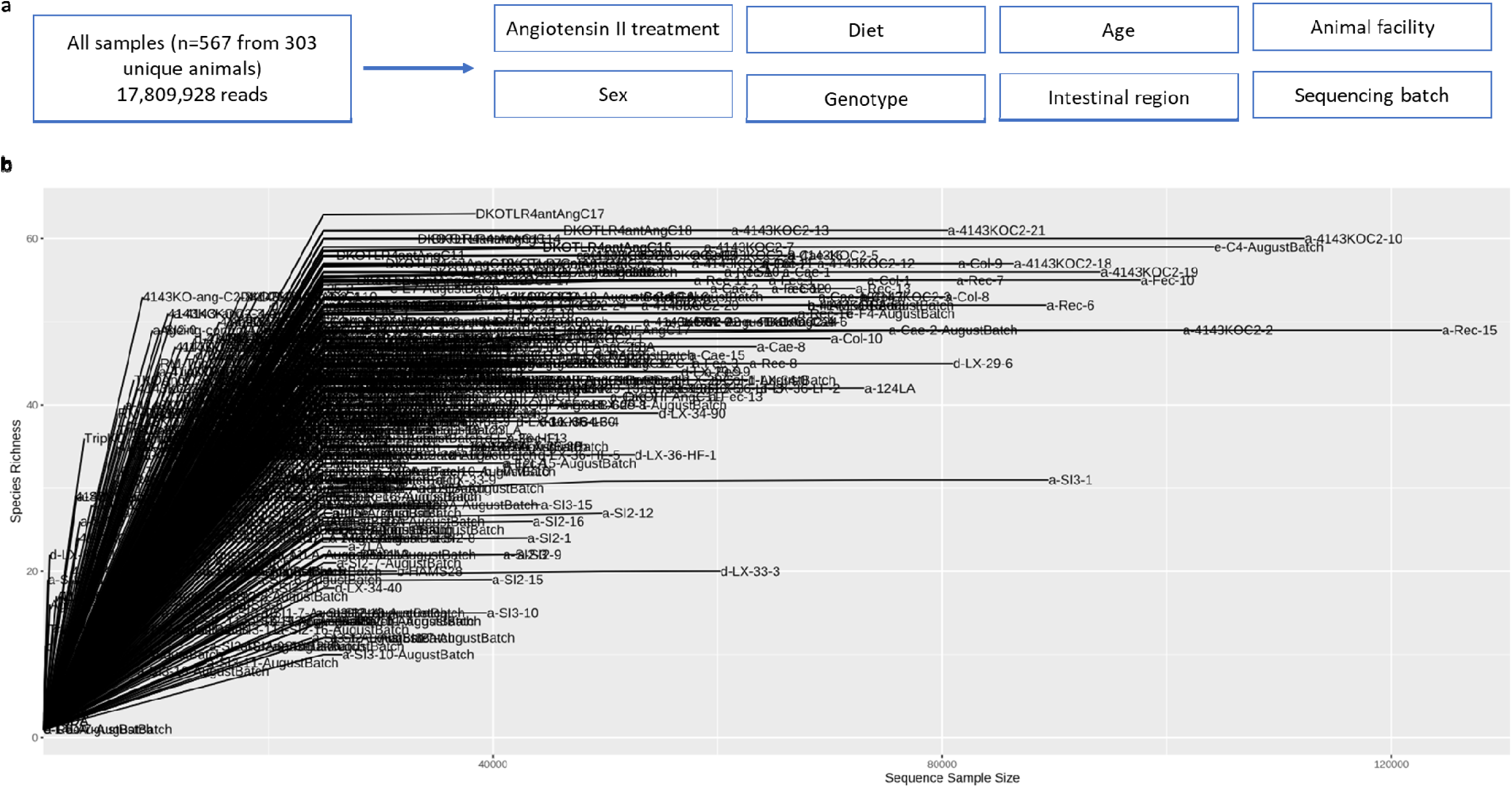
Study design and number of sequences per sample. **a**, Brief schematic of study design. 567 samples were sequenced from 303 animals, resulting in 17,809,928 reads. All samples were interrogated for various experimental factors; treatment of angiotensin II, diet, age, sex, genotype, intestinal region of sample collection site, animal facility, and sequencing batch. **b,** Number of sequences per sample.

### The impact of angiotensin II

We investigated whether high blood pressure induced by angiotensin II resulted in differences in the gut microbiota. We found significant differences in both α-(P=0.0137, Figure 2a) and β-diversity (P=0.001, Figure 2b). We identified 20 significantly differentially abundant taxa including *Clostridium leptum* (Table S2) – known to produce short-chain fatty acids (SCFAs), gut microbiota-derived metabolites shown to lower blood pressure in mice and hypertensive patients.^16,26–28^

**Figure 2.**
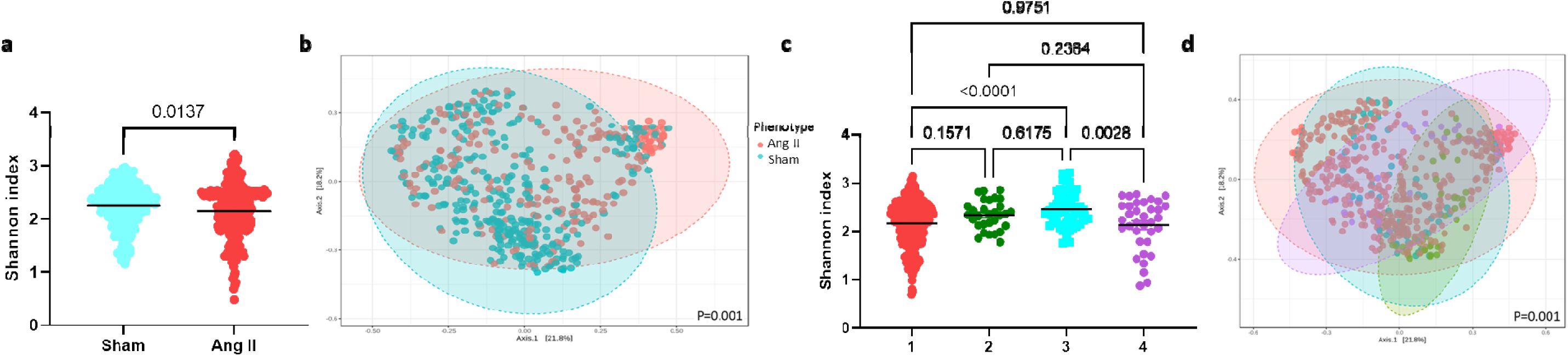
α- and β-diversity metrics of all samples (n=538), categorised according to surgery (sham versus angiotensin II) (a-b), and genotypes 1-4, with 1 being wild-type and 2-4 being knockout strains (c-d). **a, c,** Shannon index, an α-diversity metric. **b, d,** Bray-Curtis index (β-diversity metric). Significance of Shannon index (p<0.05) determined using Mann-Whitney U (two comparison groups) or Kruskal-Wallis (more than two comparison groups) tests. Significance of Bray-Curtis index determined using PERMANOVA with adjustment for multiple comparisons (q<0.05).

### The impact of the genotype

We next examined whether the mouse genotype influenced the gut bacterial metrics. We observed some significant changes in both α-diversity, particularly between genotype 1 and genotype 3 (P<0.0001), and genotypes 3 and 4 (P=0.0028,Figure 2c). There was also a significant change in β-diversity (P=0.001, Figure 2d), with some genotypes (e.g., genotype 2) being the most dissimilar across groups. Pairwise comparisons for β-diversity metrics reveal all four genotypes are significantly different to each other (Table S3). We identified 116 taxa that were different across the genotypes (Table S4), particularly in mice lacking GPR41/43/109A, which lack the 3 main receptors that sense SCFAs.

### The impact of animal house facilities

We also studied the influence of animal house facilities by using samples from three distinct facilities in Australia. We determined that animal house facility influences bacterial diversity, with differences in both Shannon index (P=0.0012-0.1103, Figure 3a) and Bray-Curtis Index (P=0.001, Figure 3b) – these show distinct clustering according to the origin of the mice. Pairwise comparisons revealed significant differences between all three facilities in Bray-Curtis Index (Table S5). We identified 95 bacteria taxa that were significantly different between animal houses (Table S6). This included common gut bacteria, such as those from the genera *Bacteroides, Alistipes, Mucispirillum* and *Azospirillum*.

**Figure 3.**
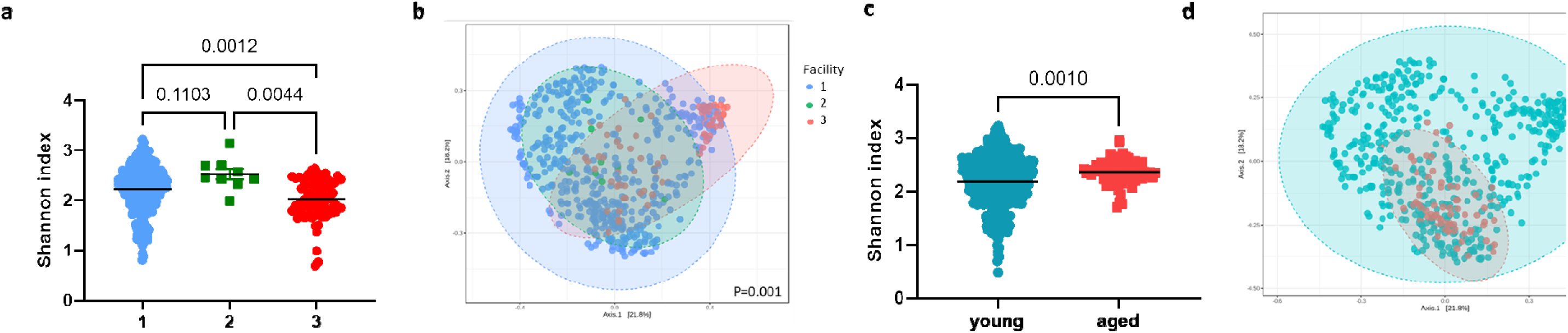
α- and β-diversity metrics of all samples (n=538), categorised according to animal facility (a-b), and age group (c-d). **a, c,** Shannon index, an α-diversity metric. **b, d,** Bray-Curtis index (β-diversity metric). Significance of Shannon index (p<0.05) determined using Mann-Whitney U (two comparison groups) or Kruskal-Wallis (more than two comparison groups) tests. Significance of Bray-Curtis index determined using PERMANOVA with adjustment for multiple comparisons (q<0.05).

### The impact of age

We categorised the samples into young (10-12 weeks) and aged (6 months) groups based on age. Although there was a large difference in sample size between young and aged mice, we found that aged mice had higher α-diversity (P<0.0001, Figure 3c) and distinct β-diversity metrics compared to younger mice (P=0.001, Figure 3d). We also identified 53 taxonomic changes between young and aged mice, with younger mice having higher prevalence of SCFA-producers such as *Ruminococcus* and *Blautia* compared to aged mice (Table S7).

### The impact of sex

There is growing interest in the representation of both male and female mice in animal studies, however, it is still unclear if there are differences in the gut microbiota driven by sex.^29^ We found that male and female mice did not differ significantly in α-diversity (P=0.054, Figure 4a), but differed in terms of β-diversity metrics (Figure 4b). Albeit not distinct in the PCoA plots, sex reached statistical significance (P=0.001), with 35 taxonomic changes between the sexes (Table S7). Male mice had lower levels of SCFA-producing bacteria including *Roseburia*, *Blautia* and *Lachnospiraceae bacterium,* relative to female mice.

**Figure 4.**
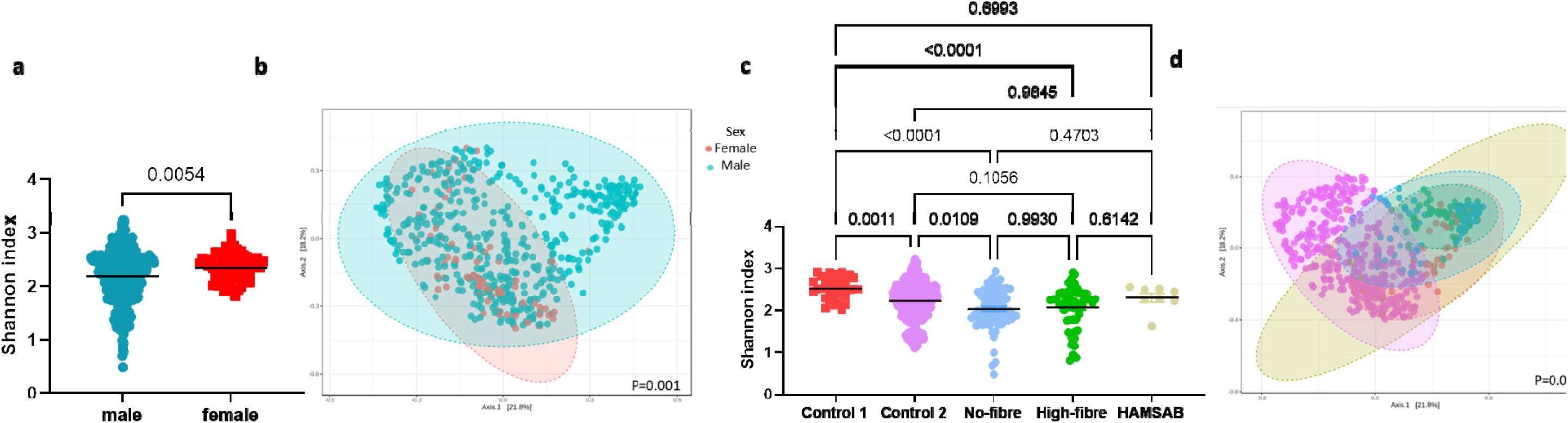
α- and β-diversity metrics of all samples (n=538), categorised according to sex (a-b), and diets (c-d). **a, c,** Shannon index, an α-diversity metric. **b, d,** Bray-Curtis index (β-diversity metric). Significance of Shannon index (p<0.05) determined using Mann-Whitney U (two comparison groups) or Kruskal-Wallis (more than two comparison groups) tests. Significance of Bray-Curtis index determined using PERMANOVA with adjustment for multiple comparisons (q<0.05).

### The impact of diet

Diet is a well-known factor that impacts the microbiome.^30^ We investigated the influence of diet using five different diets, four of which were manipulated in the same background, so the only difference was the amount and type of dietary fibre, a key prebiotic nutrient that feeds the microbiota. These diets had a clear impact on the microbiome – we found some differences in α-diversity particularly between control 1 and high-fibre diet (P<0.0001), control 1 and low-fibre diet (P<0.0001, Figure 4c), with an overall difference in β-diversity (P=0.001, Figure 4d). Pairwise comparisons for β-diversity metrics reveal all diets are mostly different to each other (Table S9). Importantly, there was a significant difference in both α- and β-diversity between the two control diets, showing the importance for control diets to be nutrient-matched. The different diets modulated 192 taxa (Table S10), with a no-fibre diet being the major driver of these changes (74 out of the 192 taxa). The no-fibre diet reduced the prevalence of bacteria known to ferment fibre and produce SCFAs, such as *Roseburia* and *Lachnospiraceae NK4A136 group*, while a high fibre diet increased the prevalence of genus *Akkermansia*.

### The impact of gastrointestinal tissue sampling

Due to differences in pH and bacterial density,^31^ samples from the large and small intestines were expected to differ. We confirmed that the small intestine had lower α-diversity (P<0.0001, Figure 5a) and different β-diversity metrics compared to the large intestine (P=0.001, Figure 5b). We also observed 72 taxa that changed between these intestinal regions (Table S11), with the small intestine having lower prevalence of 63 out of the 72 taxa.

**Figure 5.**
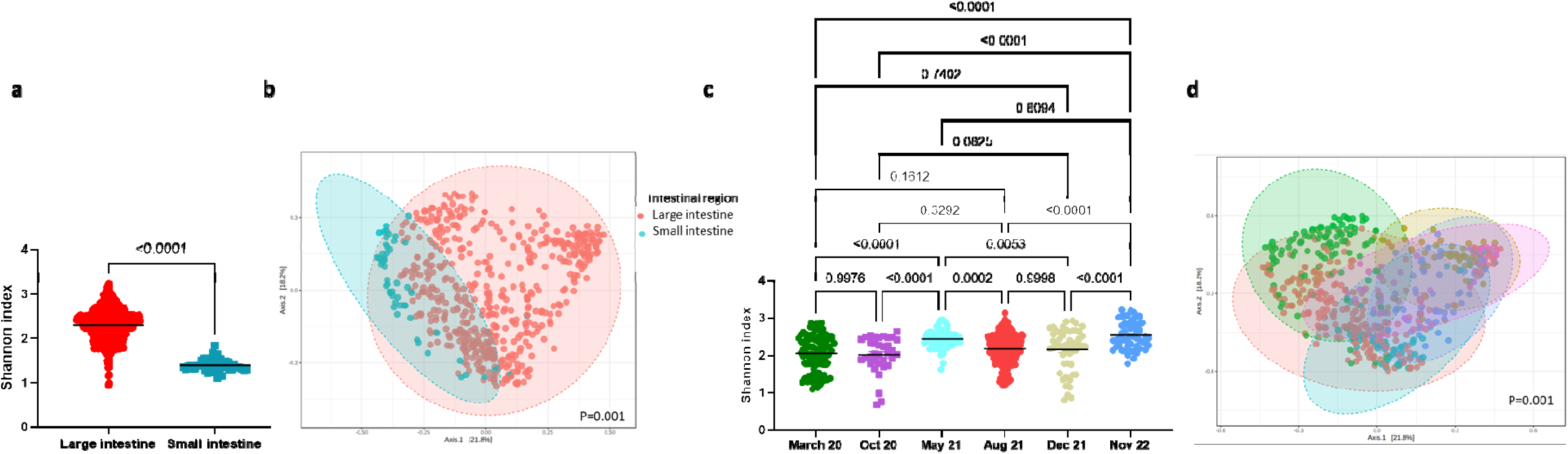
α- and β-diversity metrics of all samples (n=538), categorised according to intestinal region (a-b), and sequencing batch (c-d). **a, c,** Shannon index, an α-diversity metric. **b, d,** Bray-Curtis index (β-diversity metric). Significance of Shannon index (p<0.05) determined using Mann-Whitney U (two comparison groups) or Kruskal-Wallis (more than two comparison groups) tests. Significance of Bray-Curtis index determined using PERMANOVA with adjustment for multiple comparisons (q<0.05).

### The impact of sequencing batch effect

Sequencing batch effect is a well-known issue in which technical variations involved with sample preparation and sequencing platform results in differences in terms of bacteria identified.^32^ This poses a problem when sequenced data is combined, often observed in large-scale human studies.^33^ Samples in our study were sequenced across six batches. We found that both α- (P=0.0001-0.9998, Figure 5c) and β-diversity metrics (P=0.001, Figure 5d, Table S12) were significantly influenced by sequencing batch effect. This also resulted in 243 differential taxa across the batches (Table S13).

### Degree of variation contributed by experimental factors

We leverage a tool called VPThemAll^25^ to determine the contribution of these various factors to the variability observed in the gut microbiome. We identified that all the factors investigated in this study significantly contributed to gut microbial variations (Figure 6, Table S14). The full effects of all the factors combined explained 48.8% of variations in the gut microbiome (Table S14). Shared variances explain how much variations observed in the gut microbiome are due to each experimental factor, without correction for the other variables (Table S14). Diet had the largest influence in the shared element of the data (23.3%). The unique element, which is the variation contributed by a single factor following adjustment for all the factors, was largely contributed by the compartment from which the sample was acquired from (6.8%), followed by diet (6%), and then sequencing batch (4.7%) (Table S14). Age, genotype, and animal facility each explained ∼2% of variance in the gut microbial composition (Table S14). Sex and the treatment of angiotensin II had minimal effects on the gut microbial composition, each explaining 0.4% of the variance (Table S14).

**Figure 6.**
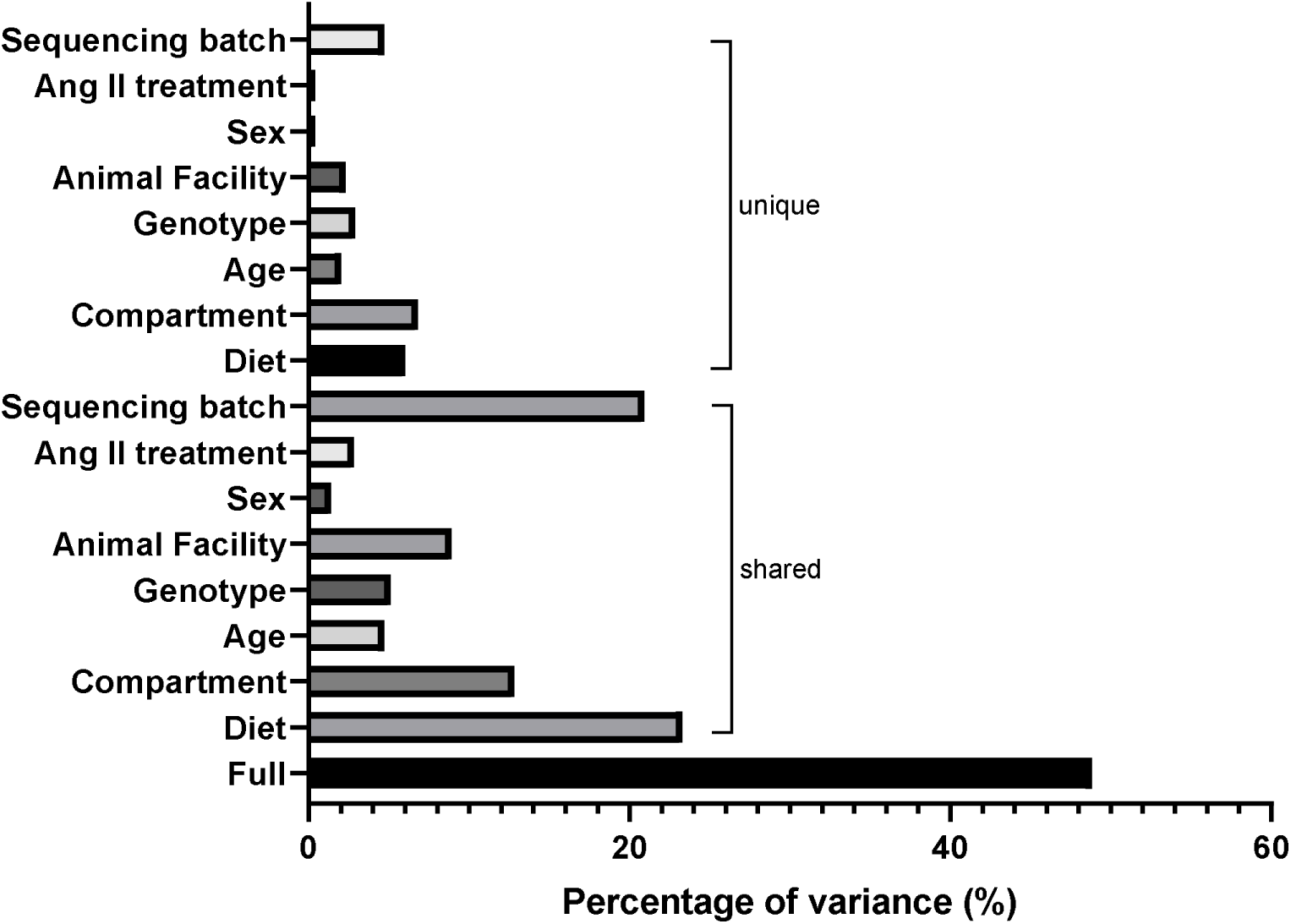
The full, shared, and unique contribution of each experimental factor to the variations in the gut microbiota composition. Full represents how much all the experimental factors in our study explain the variations observed in the gut microbiome. Shared represents how much each factor explains the variation without correcting for the other variables. Unique is how much each factor explains the variations observed, after correcting for all other variables. Refer to **Table S14** for further information.

We also performed a stepwise multiple-regression analysis to determine which were the key factors drive α-diversity. Gastrointestinal compartment (β=-0.822, CI 95%: −0.939 to −0.706, P<0.001), animal house facility (β=-0.227, CI 95%: −0.309 to −0.145, P<0.001), and age group (β=-0.132, CI 95%: 0.032 to 0.232, P=0.009) were the only factors associated with Shannon index in our cohort. Gastrointestinal compartment had the largest impact (standardised β=-0.503) relative to facility (standardised β=-0.197) and age group (standardised β=0.094).

## Discussion

We performed a retrospective study to determine the role of angiotensin II and several other experimental factors on the gut microbiome of laboratory mice. We sequenced gut microbiome samples from an heterogenous cohort of mice to introduce the influence of external factors such as diet, living environment, differences in blood pressure, genetics/genotype, sex, and age. Despite the presence of confounding factors, our study findings are in agreement with previous small-scale studies showing that angiotensin II influenced microbial diversity.^14–16,34^ Despite the influence of confounding factors, normalised data in a highly powered cohort found a few differentially abundant bacteria in common with previous studies. However, we determined that only 0.4% of the gut microbial composition differences were due to the treatment with angiotensin II, and angiotensin II was not associated with α-diversity when all factors were considered. Our findings also revealed that diet and intestinal region where samples were collected from had a large influence on the variations in the gut microbiome. Lack of appropriate control of those may lead to misleading findings that could be attributed to disease phenotypes such as angiotensin II.

The gut microbiome is often influenced by many external confounding factors which are often difficult to be controlled for. To control for these factors at the experimental level, many animals would be required, which is not feasible nor ethical. One of the approaches to overcome this issue is to validate findings across different facilities or by performing meta-analysis of multiple studies. We took advantage of the availability of samples processed by the same laboratory and team, but from across multiple facilities and cohorts with the influence of many external factors. To our knowledge, this is the largest sample size ever analysed in laboratory mouse gut microbiome (Table S1). The findings of our study highlight the potential of using a retrospective approach of a heterogenous cohort of laboratory animals as a validation method of small-scale pre-clinical animal models given that factors influencing the gut microbiome are reported.

Our study also highlights that there are many confounding factors that influence the variations observed in the gut microbiome. In addition, we were able to determine the magnitude by which these factors influence the gut microbiome – unsurprisingly, diet and the intestinal region where the sample was collected were the largest contributors to the variations observed in the gut microbiome. The findings support guidelines for gut microbiome studies in hypertension and renal function, where appropriate control for these factors has been suggested.^35,36^

In our study, we investigated the bacterial component of the gut microbiome as a proof-of-concept. However, how signatures of other members of the gut microbiome such as phages and fungi are affected by external factors remain unclear. The shared contribution of the factors we investigated only accounted for about 50% of the observed variation in the gut microbiome, meaning other factors are also at play. Other experimental factors not studied here may also impact the gut microbiome of experimental animals. This includes animal vendor,^37^ which could not be addressed in our study as all animals included were bred in-house in each of the three facilities studied. Other factors which were not considered in this study include cage mate effect prior to surgery, bedding, and despite the fact that the mice are from specific pathogen-free facilities, the presence of different pathogens can influence the gut microbiome too.^6,38,39^ How these factors interact with each other is also unclear.

We found that less than 1% of the gut microbiome was influenced by the treatment of angiotensin II. This suggests that if experimental groups are not appropriately controlled by other factors known to influence the gut microbiome, a false-positive association is likely. An example is the significant difference between the two control diets we compared. In addition, any study with sufficient sample size would detect a statistical association, but whether this is of biological significance depends on the study design.

In future studies, it would be interesting to determine the magnitude that dietary salt contributes to variations in the gut microbiome in addition to angiotensin II, as well as a meta-analysis to compare the gut microbiome across different models of hypertension. At the moment, this is a limitation of the field as many studies did not make microbiome data publicly available.

In conclusion, our large-scale retrospective study validated findings from small-scale studies that angiotensin II treatment influences the gut microbiome; however, this effect was very small relative to most of the other factors studied. Our findings also demonstrate the potential use of a heterogenous animal cohort to retrospectively investigate the role of the gut microbiome in pre-clinical disease models while being able to control for confounding variables. Controlling for these factors in future pre-clinical studies in experimental hypertension will increase the likelihood that microbiome findings are replicable and translatable. This approach could be used for other cardiovascular disease models to validate gut microbiome findings from small-scale animal studies.

## Declarations

### Ethics approval

All samples used in this study were collected for independent studies where animal ethics approval was previously received. We retrospectively reanalysed all the samples.

### Availability of data and materials

Sequencing data and metadata file are publicly available at https://doi.org/10.5281/zenodo.7935261

## Acknowledgements

We would like to acknowledge the Monash Animal Research Platform for support with animal work and the Monash Bioinformatics Platform for access to M3 servers. The authors also acknowledge the use of the services and facilities of Australian Genome Research Facility.

## Authors’ Contributions

R.R.M. designed the study, planned, and performed the data analyses, provided intellectual inputs, and wrote the manuscript. M.N., T.Z., and M.S. assisted with bioinformatics analyses, E.D., L.X., H.J., M.P., W.S., F.W., A.V., D.K., and C.R.M. provided samples from their independent studies. F.Z.M. designed the study, secured funding, provided intellectual input, and supervised the study. All authors had full access to all data in the study, revised the manuscript critically, approved the version to be published, and had the final responsibility for the decision to submit the manuscript for publication.

## References

1. Turnbaugh PJ, Bäckhed F, Fulton L, Gordon JI. Diet-Induced Obesity Is Linked to Marked but Reversible Alterations in the Mouse Distal Gut Microbiome. Cell Host & Microbe 2008;3:213–223.

2. Hufeldt MR, Nielsen DS, Vogensen FK, Midtvedt T, Hansen AK. Variation in the gut microbiota of laboratory mice is related to both genetic and environmental factors. Comp Med 2010;60:336–347.

3. Hildebrand F, Nguyen TLA, Brinkman B, Yunta RG, Cauwe B, Vandenabeele P, Liston A, Raes J. Inflammation-associated enterotypes, host genotype, cage and inter-individual effects drive gut microbiota variation in common laboratory mice. Genome Biology 2013;14:R4.

4. Rausch P, Basic M, Batra A, Bischoff SC, Blaut M, Clavel T, Gläsner J, Gopalakrishnan S, Grassl GA, Günther C, Haller D, Hirose M, Ibrahim S, Loh G, Mattner J, Nagel S, Pabst O, Schmidt F, Siegmund B, Strowig T, Volynets V, Wirtz S, Zeissig S, Zeissig Y, Bleich A, Baines JF. Analysis of factors contributing to variation in the C57BL/6J fecal microbiota across German animal facilities. International Journal of Medical Microbiology 2016;306:343–355.

5. Hilbert T, Steinhagen F, Senzig S, Cramer N, Bekeredjian-Ding I, Parcina M, Baumgarten G, Hoeft A, Frede S, Boehm O, Klaschik S. Vendor effects on murine gut microbiota influence experimental abdominal sepsis. J Surg Res 2017;211:126–136.

6. Ericsson AC, Gagliardi J, Bouhan D, Spollen WG, Givan SA, Franklin CL. The influence of caging, bedding, and diet on the composition of the microbiota in different regions of the mouse gut. Sci Rep 2018;8:4065.

7. Montonye DR, Ericsson AC, Busi SB, Lutz C, Wardwell K, Franklin CL. Acclimation and Institutionalization of the Mouse Microbiota Following Transportation. Frontiers in Microbiology 2018;9.

8. Robertson SJ, Lemire P, Maughan H, Goethel A, Turpin W, Bedrani L, Guttman DS, Croitoru K, Girardin SE, Philpott DJ. Comparison of Co-housing and Littermate Methods for Microbiota Standardization in Mouse Models. Cell Rep 2019;27:1910–1919.e1912.

9. Wolff NS, Jacobs MC, Haak BW, Roelofs JJTH, de Vos AF, Hugenholtz F, Wiersinga WJ. Vendor effects on murine gut microbiota and its influence on lipopolysaccharide-induced lung inflammation and Gram-negative pneumonia. Intensive Care Medicine Experimental 2020;8:47.

10. Singh G, Brass A, Cruickshank SM, Knight CG. Cage and maternal effects on the bacterial communities of the murine gut. Scientific Reports 2021;11:9841.

11. Guo J, Song C, Liu Y, Wu X, Dong W, Zhu H, Xiang Z, Qin C. Characteristics of gut microbiota in representative mice strains: Implications for biological research. Animal Model Exp Med 2022;5:337–349.

12. Weiss S, Xu ZZ, Peddada S, Amir A, Bittinger K, Gonzalez A, Lozupone C, Zaneveld JR, Vázquez-Baeza Y, Birmingham A, Hyde ER, Knight R. Normalization and microbial differential abundance strategies depend upon data characteristics. Microbiome 2017;5:27.

13. Jama HA, Muralitharan RR, Xu C, O’Donnell JA, Bertagnolli M, Broughton BRS, Head GA, Marques FZ. Rodent models of hypertension. Br J Pharmacol 2022;179:918–937.

14. Yang T, Santisteban MM, Rodriguez V, Li E, Ahmari N, Carvajal JM, Zadeh M, Gong M, Qi Y, Zubcevic J, Sahay B, Pepine CJ, Raizada MK, Mohamadzadeh M. Gut dysbiosis is linked to hypertension. Hypertension 2015;65:1331–1340.

15. Cheema MU, Pluznick JL. Gut Microbiota Plays a Central Role to Modulate the Plasma and Fecal Metabolomes in Response to Angiotensin II. Hypertension 2019;74:184–193.

16. Kim S, Goel R, Kumar A, Qi Y, Lobaton G, Hosaka K, Mohammed M, Handberg Eileen M, Richards Elaine M, Pepine Carl J, Raizada Mohan K. Imbalance of gut microbiome and intestinal epithelial barrier dysfunction in patients with high blood pressure. Clinical Science 2018;132:701.

17. Caporaso JG, Lauber CL, Walters WA, Berg-Lyons D, Huntley J, Fierer N, Owens SM, Betley J, Fraser L, Bauer M, Gormley N, Gilbert JA, Smith G, Knight R. Ultra-high-throughput microbial community analysis on the Illumina HiSeq and MiSeq platforms. The ISME Journal 2012;6:1621–1624.

18. Bolyen E, Rideout JR, Dillon MR, Bokulich NA, Abnet CC, Al-Ghalith GA, Alexander H, Alm EJ, Arumugam M, Asnicar F, Bai Y, Bisanz JE, Bittinger K, Brejnrod A, Brislawn CJ, Brown CT, Callahan BJ, Caraballo-Rodríguez AM, Chase J, Cope EK, Da Silva R, Diener C, Dorrestein PC, Douglas GM, Durall DM, Duvallet C, Edwardson CF, Ernst M, Estaki M, Fouquier J, Gauglitz JM, Gibbons SM, Gibson DL, Gonzalez A, Gorlick K, Guo J, Hillmann B, Holmes S, Holste H, Huttenhower C, Huttley GA, Janssen S, Jarmusch AK, Jiang L, Kaehler BD, Kang KB, Keefe CR, Keim P, Kelley ST, Knights D, Koester I, Kosciolek T, Kreps J, Langille MGI, Lee J, Ley R, Liu Y-X, Loftfield E, Lozupone C, Maher M, Marotz C, Martin BD, McDonald D, McIver LJ, Melnik AV, Metcalf JL, Morgan SC, Morton JT, Naimey AT, Navas-Molina JA, Nothias LF, Orchanian SB, Pearson T, Peoples SL, Petras D, Preuss ML, Pruesse E, Rasmussen LB, Rivers A, Robeson MS, Rosenthal P, Segata N, Shaffer M, Shiffer A, Sinha R, Song SJ, Spear JR, Swafford AD, Thompson LR, Torres PJ, Trinh P, Tripathi A, Turnbaugh PJ, Ul-Hasan S, van der Hooft JJJ, Vargas F, Vázquez-Baeza Y, Vogtmann E, von Hippel M, Walters W, Wan Y, Wang M, Warren J, Weber KC, Williamson CHD, Willis AD, Xu ZZ, Zaneveld JR, Zhang Y, Zhu Q, Knight R, Caporaso JG. Reproducible, interactive, scalable and extensible microbiome data science using QIIME 2. Nature Biotechnology 2019;37:852–857.

19. Callahan BJ, McMurdie PJ, Rosen MJ, Han AW, Johnson AJA, Holmes SP. DADA2: High-resolution sample inference from Illumina amplicon data. Nature Methods 2016;13:581–583.

20. Bokulich NA, Kaehler BD, Rideout JR, Dillon M, Bolyen E, Knight R, Huttley GA, Gregory Caporaso J. Optimizing taxonomic classification of marker-gene amplicon sequences with QIIME 2’s q2-feature-classifier plugin. Microbiome 2018;6:90.

21. Quast C, Pruesse E, Yilmaz P, Gerken J, Schweer T, Yarza P, Peplies J, Glöckner FO. The SILVA ribosomal RNA gene database project: improved data processing and web-based tools. Nucleic Acids Res 2013;41:D590–D596.

22. Chong J, Liu P, Zhou G, Xia J. Using MicrobiomeAnalyst for comprehensive statistical, functional, and meta-analysis of microbiome data. Nature Protocols 2020;15:799–821.

23. Lu Y, Zhou G, Ewald J, Pang Z, Shiri T, Xia J. MicrobiomeAnalyst 2.0: comprehensive statistical, functional and integrative analysis of microbiome data. Nucleic Acids Res 2023;51:W310–W318.

24. Mallick H, Rahnavard A, McIver LJ, Ma S, Zhang Y, Nguyen LH, Tickle TL, Weingart G, Ren B, Schwager EH, Chatterjee S, Thompson KN, Wilkinson JE, Subramanian A, Lu Y, Waldron L, Paulson JN, Franzosa EA, Bravo HC, Huttenhower C. Multivariable association discovery in population-scale meta-omics studies. PLOS Computational Biology 2021;17:e1009442.

25. Forslund SK, Chakaroun R, Zimmermann-Kogadeeva M, Markó L, Aron-Wisnewsky J, Nielsen T, Moitinho-Silva L, Schmidt TSB, Falony G, Vieira-Silva S, Adriouch S, Alves RJ, Assmann K, Bastard J-P, Birkner T, Caesar R, Chilloux J, Coelho LP, Fezeu L, Galleron N, Helft G, Isnard R, Ji B, Kuhn M, Le Chatelier E, Myridakis A, Olsson L, Pons N, Prifti E, Quinquis B, Roume H, Salem J-E, Sokolovska N, Tremaroli V, Valles-Colomer M, Lewinter C, Søndertoft NB, Pedersen HK, Hansen TH, Amouyal C, Andersson Galijatovic EA, Andreelli F, Barthelemy O, Batisse J-P, Belda E, Berland M, Bittar R, Blottière H, Bosquet F, Boubrit R, Bourron O, Camus M, Cassuto D, Ciangura C, Collet J-P, Dao M-C, Djebbar M, Doré A, Engelbrechtsen L, Fellahi S, Fromentin S, Galan P, Gauguier D, Giral P, Hartemann A, Hartmann B, Holst JJ, Hornbak M, Hoyles L, Hulot J-S, Jaqueminet S, Jørgensen NR, Julienne H, Justesen J, Kammer J, Krarup N, Kerneis M, Khemis J, Kozlowski R, Lejard V, Levenez F, Lucas-Martini L, Massey R, Martinez-Gili L, Maziers N, Medina-Stamminger J, Montalescot G, Moute S, Neves AL, Olanipekun M, Le Pavin LP, Poitou C, Pousset F, Pouzoulet L, Rodriguez-Martinez A, Rouault C, Silvain J, Svendstrup M, Swartz T, Vanduyvenboden T, Vatier C, Walther S, Gøtze JP, Køber L, Vestergaard H, Hansen T, Zucker J-D, Hercberg S, Oppert J-M, Letunic I, Nielsen J, Bäckhed F, Ehrlich SD, Dumas M-E, Raes J, Pedersen O, Clément K, Stumvoll M, Bork P, The MetaCardis C. Combinatorial, additive and dose-dependent drug– microbiome associations. Nature 2021;600:500–505.

26. Marques FZ, Nelson E, Chu P-Y, Horlock D, Fiedler A, Ziemann M, Tan Jian K, Kuruppu S, Rajapakse Niwanthi W, El-Osta A, Mackay Charles R, Kaye David M. High-Fiber Diet and Acetate Supplementation Change the Gut Microbiota and Prevent the Development of Hypertension and Heart Failure in Hypertensive Mice. Circulation 2017;135:964–977.

27. Bartolomaeus H, Balogh A, Yakoub M, Homann S, Markó L, Höges S, Tsvetkov D, Krannich A, Wundersitz S, Avery EG, Haase N, Kräker K, Hering L, Maase M, Kusche-Vihrog K, Grandoch M, Fielitz J, Kempa S, Gollasch M, Zhumadilov Z, Kozhakhmetov S, Kushugulova A, Eckardt K-U, Dechend R, Rump LC, Forslund SK, Müller DN, Stegbauer J, Wilck N. Short-Chain Fatty Acid Propionate Protects From Hypertensive Cardiovascular Damage. Circulation 2019;139:1407–1421.

28. Jama HA, Rhys-Jones D, Nakai M, Yao CK, Climie RE, Sata Y, Anderson D, Creek DJ, Head GA, Kaye DM, Mackay CR, Muir J, Marques FZ. Prebiotic intervention with HAMSAB in untreated essential hypertensive patients assessed in a phase II randomized trial. Nature Cardiovascular Research 2023;2:35–43.

29. Beale AL, Kaye DM, Marques FZ. The role of the gut microbiome in sex differences in arterial pressure. Biol Sex Differ 2019;10:22.

30. Leeming ER, Johnson AJ, Spector TD, Le Roy CI. Effect of Diet on the Gut Microbiota: Rethinking Intervention Duration. Nutrients 2019;11:2862.

31. O’Donnell JA, Zheng T, Meric G, Marques FZ. The gut microbiome and hypertension. Nature Reviews Nephrology 2023;19:153–167.

32. Lozupone CA, Stombaugh J, Gonzalez A, Ackermann G, Wendel D, Vázquez-Baeza Y, Jansson JK, Gordon JI, Knight R. Meta-analyses of studies of the human microbiota. Genome Res 2013;23:1704–1714.

33. de Goffau MC, Charnock-Jones DS, Smith GCS, Parkhill J. Batch effects account for the main findings of an in utero human intestinal bacterial colonization study. Microbiome 2021;9:6.

34. Avery EG, Bartolomaeus H, Rauch A, Chen CY, N’Diaye G, Löber U, Bartolomaeus TUP, Fritsche-Guenther R, Rodrigues AF, Yarritu A, Zhong C, Fei L, Tsvetkov D, Todiras M, Park JK, Markó L, Maifeld A, Patzak A, Bader M, Kempa S, Kirwan JA, Forslund SK, Müller DN, Wilck N. Quantifying the impact of gut microbiota on inflammation and hypertensive organ damage. Cardiovascular Research 2022.

35. Muralitharan RR, Snelson M, Meric G, Coughlan MT, Marques FZ. Guidelines for Microbiome Studies in Renal Physiology. Am J Physiol Renal Physiol 2023.

36. Marques FZ, Jama HA, Tsyganov K, Gill PA, Rhys-Jones D, Muralitharan RR, Muir JG, Holmes AJ, Mackay CR. Guidelines for Transparency on Gut Microbiome Studies in Essential and Experimental Hypertension. Hypertension 2019;74:1279–1293.

37. Rasmussen TS, de Vries L, Kot W, Hansen LH, Castro-Mejía JL, Vogensen FK, Hansen AK, Nielsen DS. Mouse Vendor Influence on the Bacterial and Viral Gut Composition Exceeds the Effect of Diet. Viruses 2019;11.

38. Ridaura VK, Faith JJ, Rey FE, Cheng J, Duncan AE, Kau AL, Griffin NW, Lombard V, Henrissat B, Bain JR, Muehlbauer MJ, Ilkayeva O, Semenkovich CF, Funai K, Hayashi DK, Lyle BJ, Martini MC, Ursell LK, Clemente JC, Van Treuren W, Walters WA, Knight R, Newgard CB, Heath AC, Gordon JI. Gut microbiota from twins discordant for obesity modulate metabolism in mice. Science 2013;341:1241214.

39. Gregor A, Fragner L, Trajanoski S, Li W, Sun X, Weckwerth W, König J, Duszka K. Cage bedding modifies metabolic and gut microbiota profiles in mouse studies applying dietary restriction. Scientific Reports 2020;10:20835.

