## Supplementary material for "The Influence of Angiotensin II on the Gut Microbiome of Mice: Findings from a Retrospective Study": Online supplemental tables and figures

[1]Hypertension Research Laboratory, School of Biological Sciences, Faculty of Science, Monash University, Melbourne, Australia; [2]Institute for Medical Research, Ministry of Health Malaysia, Kuala Lumpur, Malaysia; [3]Victorian Heart Institute, Monash University, Melbourne, Australia.; [4]Heart Failure Research Group, Baker Heart and Diabetes Institute, Melbourne, Australia; [5]Department of Microbiology, Anatomy, Physiology, La Trobe University, Melbourne, Australia; [6]Department of Cardiology, Alfred Hospital, Melbourne, Australia; [7]Central Clinical School, Faculty of Medicine Nursing and Health Sciences, Monash University, Melbourne, Australia;[8]Infection and Immunity Program, Monash Biodiscovery Institute, Monash University, Melbourne, Australia; [9]Department of Biochemistry, Monash University, Melbourne, Australia; [10]School of Pharmaceutical Sciences, Shandong Analysis and Test Center, Qilu University of Technology (Shandong Academy of Sciences), Jinan, 250014, China;

***Correspondence to**: A/Prof Francine Marques. Hypertension Research Laboratory, School of Biological Sciences, Monash University, 25 Rainforest Walk, Clayton, Victoria 3800, Australia. P: +61-03-99056958. E:

### Supplementary Tables

**Table S1.** Summary of previous studies that investigated the role of experimental factors that influence the gut bacterial diversity metrics in laboratory mice. Search performed on PubMed on 9$^{th}$ March 2023 using the following keywords: 'alpha diversity', 'beta diversity', 'experimental factors', 'laboratory', 'mice', 'animals'.

| Study | Factor(s) investigated | Total sample size | Findings |
|---|---|---|---|
| Turnbaugh et al., 2008 | Diet | 10 | Conventionalised germ-free mice fed two different diets had significant difference in β-diversity |
| Hufeldt et al., 2010 | Mice vendor | 33 | C57BL/6 mice from 2 different vendors had different caecal microbiota composition |
| | Different rooms in same breeding center | 33 | C57BL/6 mice from 1 vendor bred in two different rooms in the same breeding centre had similar microbiota composition |
| | Sex | 33 | No difference between male and female mice from same vendor housed in the same room but different cages |
| Hildebrand et al., 2013 | Genetic effects | 50 | Genetic effects influenced up to 15.65 to 18.62% of variations at lower than phylum levels among 4 genotypes |
| | Cage effects | 50 | Up to 30% of variations in the gut microbiome were explained by cage effects |
| Rausch et al., 2016 | Housing type | 94 | Difference in α- and β-diversity in mice housed in IVC compared to others |
| | Chow treatment | 104 | Difference in α- and β-diversity in autoclaved food compared to irradiated and untreated |
| | Chow provider | 104 | No difference in α-diversity between two chow providers, but different β-diversity |
| Hilbert et al., 2017 | Vendor effects | 60 | C57BL/6 from three vendors had distinct gut microbial composition |

| | | | |
|---|---|---|---|
| Ericsson et al., 2018 | Cage ventilation and bedding | 144 | Cage ventilation and bedding and the interaction, had a distinguishable effect on the caecal microbiota but not in the faecal or small intestine samples |
| Montonye et al., 2018 | Transport | 16 | Significant difference in α- and β-diversity pre- and post-arrival |
| | Animal facility | 8 | The α- and β-diversity of mice pre-shipping is significantly different to microbiota upon arrival at facilities and between facilities |
| Robertson et al., 2019 | Mice vendor | 8 | Mice from 2 vendors had no difference in α-diversity but significantly different β-diversity. Samples from pellet, colon, ileum |
| | Littermate effect and co-housing effect | 28 | Mice that were co-housed had similar gut microbiome to each other but also their non-co-housed siblings (n=12) in the colon and the ileum but not in the faecal pellet. |
| Wolff et al., 2020 | Vendor effects | 24 | Significant difference in β-diversity of genetically identical C57BL/6 from three different vendors |
| Singh et al., 2021 | Cage and maternal effects | 30 | The gut microbiome of F2 littermates of different genotypes (n=10 per group) from heterozygous parents (mdr1a-/+) clustered according to their maternal origin and cages. |
| Guo et al., 2022 | Mouse genotypes | 110 | Significant difference in α- and β-diversity of 7 background genotypes of C57BL/6 and BALB/c mice |

**Table S2**. Differences in gut microbiome taxa between angiotensin II treatment relative to sham, showing taxa with q<0.05.

| **Taxa** | **Log2FC** | **St.Error** | **P-value** | **FDR** |
|---|---|---|---|---|
| Sham/Angiotensin II | | | | |
| p__Firmicutes | 0.422 | 0.145 | 0.00385 | 0.0101 |
| c__Gammaproteobacteria | 0.896 | 0.248 | 3.22E-04 | 0.00121 |
| o__Burkholderiales | 0.896 | 0.248 | 3.22E-04 | 0.0013 |
| o__Erysipelotrichales | 0.82 | 0.267 | 0.00225 | 0.00698 |
| o__Oscillospirales | 0.599 | 0.206 | 0.00377 | 0.0112 |
| f__Sutterellaceae | 0.896 | 0.248 | 3.22E-04 | 0.00131 |
| f__Erysipelotrichaceae | 0.82 | 0.267 | 0.00225 | 0.00719 |
| f__Oscillospiraceae | 0.596 | 0.207 | 0.00418 | 0.0126 |
| g__Oscillibacter | 0.985 | 0.241 | 5.15E-05 | 2.52E-04 |
| g__Parasutterella | 0.896 | 0.248 | 3.22E-04 | 0.00127 |
| g__uncultured | 0.537 | 0.19 | 0.00494 | 0.0141 |
| g__Colidextribacter | 0.653 | 0.237 | 0.00605 | 0.0167 |

| | | | | |
|---|---|---|---|---|
| g__Lachnoclostridium | 0.641 | 0.235 | 0.00664 | 0.018 |
| g__ASF356 | 0.619 | 0.261 | 0.0179 | 0.042 |
| g__Bilophila | 0.536 | 0.228 | 0.019 | 0.0443 |
| Not_Assigned | 0.369 | 0.159 | 0.021 | 0.0487 |
| g__Desulfovibrio | 0.543 | 0.236 | 0.0216 | 0.0499 |
| unidentified | 0.68 | 0.224 | 0.00246 | 0.00716 |
| Clostridium_leptum | 0.693 | 0.244 | 0.00466 | 0.0128 |
| uncultured_bacterium | 0.192 | 0.084 | 0.0227 | 0.0468 |

**Legend:** d (domain), p (phylum), c (class), o (order), f (family), g (genus), standard error (St. Error), false-discovery rate adjusted p-value (FDR)

**Table S3**. Pairwise comparisons between the genotypes for Bray-Curtis index (a metric of β-diversity).

| **Comparisons** | **F-value** | **R-squared** | **P-value** | **FDR** |
|---|---|---|---|---|
| Genotype 3 vs Genotype 1 | 5.266 | 0.010422 | 0.001 | 0.001 |
| Genotype 3 vs Genotype 4 | 9.2001 | 0.087454 | 0.001 | 0.001 |
| Genotype 3 vs Genotype 2 | 15.509 | 0.14561 | 0.001 | 0.001 |
| Genotype 1 vs Genotype 4 | 5.3561 | 0.01122 | 0.001 | 0.001 |
| Genotype 1 vs Genotype 2 | 15.844 | 0.032814 | 0.001 | 0.001 |
| Genotype 4 vs Genotype 2 | 13.851 | 0.18023 | 0.001 | 0.001 |

**Table S4**. Differences in gut microbiome taxa across genotypes relative to wild-type mice (Genotype #1) (showing taxa with q<0.05).

| Name | Log2FC | St.Error | P-value | FDR |
|---|---|---|---|---|
| WT/GPR41-43-109aKO | | | | |
| p__Bacteroidota | 0.664 | 0.125 | 1.48E-07 | 1.17E-06 |
| p__Verrucomicrobiota | -3.27 | 0.714 | 5.75E-06 | 3.15E-05 |
| p__Deferribacterota | 2.47 | 0.694 | 4.15E-04 | 0.00141 |
| p__Firmicutes | -0.966 | 0.294 | 0.00108 | 0.00324 |
| c__Bacteroidia | 0.664 | 0.125 | 1.48E-07 | 1.41E-06 |
| c__Bacilli | -2.93 | 0.619 | 2.74E-06 | 1.77E-05 |
| c__Verrucomicrobiae | -3.27 | 0.714 | 5.75E-06 | 3.32E-05 |
| c__Deferribacteres | 2.47 | 0.694 | 4.15E-04 | 0.00146 |
| o__Bacteroidales | 0.664 | 0.125 | 1.48E-07 | 1.25E-06 |
| o__Verrucomicrobiales | -3.27 | 0.714 | 5.75E-06 | 3.61E-05 |
| o__Acholeplasmatales | -3.7 | 0.872 | 2.59E-05 | 1.40E-04 |
| o__Clostridia_vadinBB60_group | -1.81 | 0.496 | 2.87E-04 | 0.00117 |

| | | | | |
|---|---|---|---|---|
| o__Deferribacterales | 2.47 | 0.694 | 4.15E-04 | 0.00165 |
| o__Erysipelotrichales | -1.75 | 0.54 | 0.00123 | 0.00405 |
| o__Lachnospirales | -0.986 | 0.399 | 0.0137 | 0.0353 |
| f__Rikenellaceae | 4.07 | 0.522 | 3.24E-14 | 7.14E-13 |
| f__Eubacterium_coprostanoligenes_group | 3.23 | 0.433 | 3.58E-13 | 6.44E-12 |
| f__Bacteroidaceae | 1.5 | 0.302 | 9.42E-07 | 7.18E-06 |
| f__Akkermansiaceae | -3.27 | 0.714 | 5.75E-06 | 3.67E-05 |
| f__Acholeplasmataceae | -3.7 | 0.872 | 2.59E-05 | 1.43E-04 |
| f__Clostridia_vadinBB60_group | -1.81 | 0.496 | 2.87E-04 | 0.00118 |
| f__Deferribacteraceae | 2.47 | 0.694 | 4.15E-04 | 0.00164 |
| f__Erysipelotrichaceae | -1.75 | 0.54 | 0.00123 | 0.00413 |
| f__Lachnospiraceae | -0.986 | 0.399 | 0.0137 | 0.0351 |
| g__Muribaculum | -5.48 | 0.516 | 4.44E-24 | 1.68E-22 |
| g__Dubosiella | -3.06 | 0.365 | 4.45E-16 | 1.14E-14 |
| g__Alistipes | 4.07 | 0.522 | 3.24E-14 | 7.30E-13 |
| g__Eubacterium_coprostanoligenes_group | 3.23 | 0.433 | 3.58E-13 | 7.36E-12 |

| | | | | |
|---|---|---|---|---|
| g__Ileibacterium | -2.92 | 0.557 | 2.34E-07 | 1.89E-06 |
| g__Eubacterium_xylanophilum_group | 2.25 | 0.451 | 8.14E-07 | 6.01E-06 |
| g__Bacteroides | 1.5 | 0.302 | 9.42E-07 | 6.85E-06 |
| g__GCA_900066575 | -2.54 | 0.514 | 1.01E-06 | 7.27E-06 |
| g__Akkermansia | -3.27 | 0.714 | 5.75E-06 | 3.51E-05 |
| g__Anaerotruncus | 2.56 | 0.566 | 7.48E-06 | 4.49E-05 |
| g__Tuzzerella | -1.99 | 0.466 | 2.23E-05 | 1.15E-04 |
| g__Anaeroplasma | -3.7 | 0.872 | 2.59E-05 | 1.31E-04 |
| g__Intestinimonas | 2.27 | 0.546 | 3.71E-05 | 1.84E-04 |
| g__Desulfovibrio | 1.82 | 0.477 | 1.46E-04 | 6.03E-04 |
| g__Bilophila | -1.69 | 0.461 | 2.59E-04 | 0.00104 |
| g__Clostridia_vadinBB60_group | -1.81 | 0.496 | 2.87E-04 | 0.00112 |
| g__Mucispirillum | 2.47 | 0.694 | 4.15E-04 | 0.00155 |
| g__Lachnospiraceae_UCG_006 | -1.81 | 0.512 | 4.28E-04 | 0.00159 |
| g__Dorea | 1.55 | 0.442 | 4.74E-04 | 0.00172 |
| g__Lachnospiraceae_NK4A136_group | -1.36 | 0.484 | 0.00509 | 0.014 |

| | | | | |
|---|---|---|---|---|
| uncultured_Barnesiella | 6.87 | 0.481 | 1.39E-39 | 9.17E-38 |
| Dubosiella_newyorkensis | -3.06 | 0.365 | 4.45E-16 | 7.34E-15 |
| uncultured_Bacteroidales | 2.09 | 0.297 | 6.14E-12 | 6.75E-11 |
| Clostridium_leptum | -2.66 | 0.493 | 9.58E-08 | 6.33E-07 |
| Ileibacterium_valens | -2.92 | 0.557 | 2.34E-07 | 1.40E-06 |
| mouse_gut | 2.25 | 0.483 | 3.87E-06 | 1.92E-05 |
| unidentified | -1.27 | 0.452 | 0.00525 | 0.0128 |
| WT/GPR41-43KO | | | | |
| p__Deferribacterota | 4.21 | 0.49 | 9.32E-17 | 2.35E-15 |
| p__Verrucomicrobiota | -2.49 | 0.504 | 1.07E-06 | 7.52E-06 |
| p__Proteobacteria | -1.21 | 0.357 | 7.17E-04 | 0.00232 |
| c__Deferribacteres | 4.21 | 0.49 | 9.32E-17 | 3.02E-15 |
| c__Verrucomicrobiae | -2.49 | 0.504 | 1.07E-06 | 7.91E-06 |
| c__Alphaproteobacteria | -1.6 | 0.483 | 0.00101 | 0.00326 |
| o__Deferribacterales | 4.21 | 0.49 | 9.32E-17 | 2.80E-15 |
| o__Verrucomicrobiales | -2.49 | 0.504 | 1.07E-06 | 8.05E-06 |

| | | | | |
|---|---|---|---|---|
| o__Clostridia_vadinBB60_group | -1.26 | 0.351 | 3.71E-04 | 0.00152 |
| o__Rhodospirillales | -1.6 | 0.483 | 0.00101 | 0.00357 |
| o__Peptococcales | 0.762 | 0.288 | 0.00846 | 0.0246 |
| f__Eubacterium_coprostanoligenes_group | 3.55 | 0.306 | 5.24E-28 | 1.73E-26 |
| f__Deferribacteraceae | 4.21 | 0.49 | 9.32E-17 | 2.17E-15 |
| f__Akkermansiaceae | -2.49 | 0.504 | 1.07E-06 | 8.18E-06 |
| f__Clostridia_vadinBB60_group | -1.26 | 0.351 | 3.71E-04 | 0.00155 |
| f__Tannerellaceae | -0.98 | 0.292 | 8.32E-04 | 0.00314 |
| f__uncultured | -1.6 | 0.483 | 0.00101 | 0.00366 |
| f__Peptococcaceae | 0.762 | 0.288 | 0.00846 | 0.0245 |
| g__Eubacterium_coprostanoligenes_group | 3.55 | 0.306 | 5.24E-28 | 2.36E-26 |
| g__Mucispirillum | 4.21 | 0.49 | 9.32E-17 | 2.49E-15 |
| g__Dubosiella | -1.95 | 0.258 | 1.60E-13 | 3.39E-12 |
| g__Akkermansia | -2.49 | 0.504 | 1.07E-06 | 8.40E-06 |
| g__Faecalibaculum | 1.35 | 0.368 | 2.67E-04 | 0.00117 |
| g__Clostridia_vadinBB60_group | -1.26 | 0.351 | 3.71E-04 | 0.00153 |

| | | | | |
|---|---|---|---|---|
| g__Muribaculum | -1.24 | 0.364 | 7.08E-04 | 0.00273 |
| g__A2 | 1.33 | 0.415 | 0.00136 | 0.00462 |
| g__Bilophila | -0.912 | 0.325 | 0.00523 | 0.0154 |
| g__Eubacterium_xylanophilum_group | -0.85 | 0.318 | 0.00777 | 0.0216 |
| g__Tuzzerella | -0.806 | 0.329 | 0.0146 | 0.0372 |
| Dubosiella_newyorkensis | -1.95 | 0.258 | 1.60E-13 | 2.64E-12 |
| unidentified | -1.3 | 0.319 | 5.54E-05 | 2.39E-04 |
| uncultured_bacterium | 0.357 | 0.12 | 0.00305 | 0.00863 |
| Not_Assigned | -0.226 | 0.0886 | 0.0109 | 0.0247 |
| Lachnospiraceae_bacterium | -0.942 | 0.416 | 0.0241 | 0.0478 |
| WT/GPR65KO | | | | |
| p__Verrucomicrobiota | -3.77 | 0.659 | 1.66E-08 | 1.61E-07 |
| c__Verrucomicrobiae | -3.77 | 0.659 | 1.66E-08 | 1.80E-07 |
| c__Alphaproteobacteria | 1.98 | 0.632 | 0.00184 | 0.00575 |
| o__Verrucomicrobiales | -3.77 | 0.659 | 1.66E-08 | 1.66E-07 |
| o__Clostridia_vadinBB60_group | 2.37 | 0.458 | 3.12E-07 | 2.41E-06 |

| | | | | |
|---|---|---|---|---|
| o__Clostridia_UCG_014 | -2 | 0.527 | 1.66E-04 | 7.01E-04 |
| o__Rhodospirillales | 1.98 | 0.632 | 0.00184 | 0.00607 |
| f__Akkermansiaceae | -3.77 | 0.659 | 1.66E-08 | 1.61E-07 |
| f__Eubacterium_coprostanoligenes_group | -2.1 | 0.4 | 2.22E-07 | 1.79E-06 |
| f__Clostridia_vadinBB60_group | 2.37 | 0.458 | 3.12E-07 | 2.42E-06 |
| f__Clostridia_UCG_014 | -2 | 0.527 | 1.66E-04 | 7.15E-04 |
| f__uncultured | 1.98 | 0.632 | 0.00184 | 0.00614 |
| g__Akkermansia | -3.77 | 0.659 | 1.66E-08 | 1.87E-07 |
| g__Eubacterium_coprostanoligenes_group | -2.1 | 0.4 | 2.22E-07 | 1.88E-06 |
| g__Clostridia_vadinBB60_group | 2.37 | 0.458 | 3.12E-07 | 2.58E-06 |
| g__GCA_900066575 | -2.28 | 0.475 | 1.96E-06 | 1.41E-05 |
| g__Faecalibaculum | 2.2 | 0.481 | 5.81E-06 | 3.80E-05 |
| g__Ileibacterium | -2.04 | 0.514 | 8.48E-05 | 4.02E-04 |
| g__Clostridia_UCG_014 | -2 | 0.527 | 1.66E-04 | 7.29E-04 |
| g__Lachnoclostridium | -1.65 | 0.439 | 1.84E-04 | 8.05E-04 |
| g__Intestinimonas | -1.69 | 0.503 | 8.64E-04 | 0.00314 |

| | | | | |
|---|---|---|---|---|
| g__ASF356 | 1.51 | 0.486 | 0.00196 | 0.00629 |
| g__Muribaculum | -1.29 | 0.476 | 0.00679 | 0.0189 |
| g__Lachnospiraceae_FCS020_group | 1.38 | 0.509 | 0.007 | 0.0193 |
| g__Tuzzerella | -1.12 | 0.43 | 0.00925 | 0.0245 |
| g__Eubacterium_xylanophilum_group | -1.03 | 0.416 | 0.0138 | 0.0347 |
| g__Incertae_Sedis | -0.94 | 0.396 | 0.0178 | 0.0427 |
| Ileibacterium_valens | -2.04 | 0.514 | 8.48E-05 | 3.91E-04 |
| mouse_gut | 1.26 | 0.446 | 0.00486 | 0.0132 |
| uncultured_Barnesiella | -1.2 | 0.444 | 0.007 | 0.0178 |
| uncultured_Bacteroidales | 0.641 | 0.274 | 0.0197 | 0.0408 |

**Legend:** d (domain), p (phylum), c (class), o (order), f (family), g (genus), standard error (St. Error), false-discovery rate adjusted p-value (FDR)

**Table S5**. Pairwise comparisons between the facilities for Bray-Curtis index.

| **Comparisons** | **F-value** | **R-squared** | **P-value** | **FDR** |
|---|---|---|---|---|
| Facility 1 vs Facility 3 | 3.4034 | 0.006884 | 0.009 | 0.009 |
| Facility 1 vs Facility 2 | 43.06 | 0.071999 | 0.001 | 0.0015 |
| Facility 3 vs Facility 2 | 6.4507 | 0.07293 | 0.001 | 0.0015 |

**Table S6**. Differences in gut microbiome taxa across animal house facilities relative to animal facility #1 (showing taxa with q<0.05).

| Name | Log2FC | St.Error | P-value | FDR |
|---|---|---|---|---|
| Facility 1/Facility 3 | | | | |
| p__Deferribacterota | -5.15 | 0.886 | 1.04E-08 | 1.09E-07 |
| p__Proteobacteria | 2.48 | 0.645 | 1.32E-04 | 4.77E-04 |
| p__Bacteroidota | 0.492 | 0.159 | 0.00207 | 0.00556 |
| c__Deferribacteres | -5.15 | 0.886 | 1.04E-08 | 1.20E-07 |
| c__Gammaproteobacteria | 3.22 | 0.639 | 6.26E-07 | 5.34E-06 |
| c__Bacteroidia | 0.492 | 0.159 | 0.00207 | 0.006 |
| c__Bacilli | -1.97 | 0.791 | 0.0132 | 0.0344 |
| c__Clostridia | -1.12 | 0.451 | 0.0137 | 0.0346 |
| o__Deferribacterales | -5.15 | 0.886 | 1.04E-08 | 1.17E-07 |
| o__Burkholderiales | 3.22 | 0.639 | 6.26E-07 | 4.97E-06 |
| o__Clostridia_UCG_014 | 2.51 | 0.729 | 6.20E-04 | 0.00233 |
| o__Bacteroidales | 0.492 | 0.159 | 0.00207 | 0.00651 |
| o__Lactobacillales | -1.9 | 0.695 | 0.00643 | 0.0187 |
| o__Acholeplasmatales | -2.87 | 1.11 | 0.0101 | 0.0278 |
| o__Lachnospirales | -1.29 | 0.509 | 0.0114 | 0.0307 |
| o__Peptococcales | 1.32 | 0.521 | 0.0113 | 0.0307 |
| o__Erysipelotrichales | -1.71 | 0.689 | 0.0135 | 0.0359 |
| f__Deferribacteraceae | -5.15 | 0.886 | 1.04E-08 | 1.08E-07 |
| f__Sutterellaceae | 3.22 | 0.639 | 6.26E-07 | 4.86E-06 |
| f__Butyricicoccaceae | 2.32 | 0.534 | 1.69E-05 | 9.96E-05 |
| f__Clostridia_UCG_014 | 2.51 | 0.729 | 6.20E-04 | 0.00236 |
| f__Lactobacillaceae | -2.29 | 0.706 | 0.00125 | 0.00416 |
| f__Ruminococcaceae | -1.77 | 0.582 | 0.00254 | 0.00806 |
| f__Muribaculaceae | 1.02 | 0.367 | 0.00585 | 0.0168 |
| f__Acholeplasmataceae | -2.87 | 1.11 | 0.0101 | 0.0276 |

| | | | | |
|---|---|---|---|---|
| f__Lachnospiraceae | -1.29 | 0.509 | 0.0114 | 0.0306 |
| f__Peptococcaceae | 1.32 | 0.521 | 0.0113 | 0.0306 |
| f__Rikenellaceae | 1.66 | 0.666 | 0.0129 | 0.0344 |
| f__Bacteroidaceae | 0.953 | 0.385 | 0.0137 | 0.0355 |
| f__Erysipelotrichaceae | -1.71 | 0.689 | 0.0135 | 0.0355 |
| g__Mucispirillum | -5.15 | 0.886 | 1.04E-08 | 1.22E-07 |
| g__Intestinimonas | 3.75 | 0.696 | 1.06E-07 | 1.02E-06 |
| g__Parasutterella | 3.22 | 0.639 | 6.26E-07 | 4.95E-06 |
| g__ASF356 | -3.27 | 0.672 | 1.55E-06 | 1.12E-05 |
| Not_Assigned | -1.82 | 0.411 | 1.15E-05 | 6.89E-05 |
| g__Roseburia | -3.38 | 0.768 | 1.29E-05 | 7.61E-05 |
| g__UCG_009 | 2.32 | 0.534 | 1.69E-05 | 9.34E-05 |
| g__Blautia | -3.06 | 0.753 | 5.40E-05 | 2.63E-04 |
| g__A2 | -2.98 | 0.749 | 7.97E-05 | 3.70E-04 |
| g__Anaerotruncus | -2.59 | 0.723 | 3.70E-04 | 0.00144 |
| g__Eubacterium_xylanophilum_group | 2.03 | 0.575 | 4.58E-04 | 0.00173 |
| g__Clostridia_UCG_014 | 2.51 | 0.729 | 6.20E-04 | 0.00224 |
| g__Lactobacillus | -2.29 | 0.706 | 0.00125 | 0.00405 |
| g__Lachnospiraceae_FCS020_group | -2.2 | 0.704 | 0.0019 | 0.00591 |
| g__uncultured | -1.41 | 0.491 | 0.00416 | 0.0123 |
| g__Lachnoclostridium | -1.73 | 0.607 | 0.00468 | 0.0135 |
| g__Muribaculaceae | 1.03 | 0.367 | 0.00511 | 0.0145 |
| g__Lachnospiraceae_UCG_001 | -2.46 | 0.915 | 0.00729 | 0.0195 |
| g__Anaeroplasma | -2.87 | 1.11 | 0.0101 | 0.0258 |
| g__Alistipes | 1.66 | 0.666 | 0.0129 | 0.0323 |
| g__Bacteroides | 0.953 | 0.385 | 0.0137 | 0.034 |
| Clostridium_leptum | -3.79 | 0.629 | 3.11E-09 | 2.93E-08 |
| uncultured_Barnesiella | 3.08 | 0.614 | 7.08E-07 | 4.25E-06 |
| Lachnospiraceae_bacterium | -3.06 | 0.753 | 5.40E-05 | 2.37E-04 |
| mouse_gut | -2.13 | 0.617 | 6.10E-04 | 0.002 |

| | | | | |
|---|---|---|---|---|
| unidentified | -1.63 | 0.577 | 0.0048 | 0.013 |
| Facility 1/Facility 2 | | | | |
| p__Bacteroidota | 0.988 | 0.193 | 4.10E-07 | 3.04E-06 |
| p__Deferribacterota | 4.75 | 1.07 | 1.19E-05 | 5.74E-05 |
| p__Verrucomicrobiota | 3.25 | 1.1 | 0.00341 | 0.00914 |
| p__Firmicutes | -1.26 | 0.455 | 0.00575 | 0.0145 |
| c__Bacteroidia | 0.988 | 0.193 | 4.10E-07 | 3.69E-06 |
| c__Bacilli | -4.42 | 0.958 | 4.96E-06 | 3.09E-05 |
| c__Deferribacteres | 4.75 | 1.07 | 1.19E-05 | 6.40E-05 |
| c__Verrucomicrobiae | 3.25 | 1.1 | 0.00341 | 0.00952 |
| o__Lactobacillales | -4.4 | 0.842 | 2.51E-07 | 2.12E-06 |
| o__Bacteroidales | 0.988 | 0.193 | 4.10E-07 | 3.26E-06 |
| o__Erysipelotrichales | -3.77 | 0.835 | 7.92E-06 | 4.75E-05 |
| o__Deferribacterales | 4.75 | 1.07 | 1.19E-05 | 6.81E-05 |
| o__Acholeplasmatales | -4.11 | 1.35 | 0.00243 | 0.00744 |
| o__Verrucomicrobiales | 3.25 | 1.1 | 0.00341 | 0.01 |
| f__Lactobacillaceae | -4.28 | 0.855 | 7.64E-07 | 6.05E-06 |
| f__Eubacterium_coprostanoligenes_group | 3.16 | 0.67 | 3.17E-06 | 2.20E-05 |
| f__Erysipelotrichaceae | -3.77 | 0.835 | 7.92E-06 | 4.90E-05 |
| f__Deferribacteraceae | 4.75 | 1.07 | 1.19E-05 | 7.11E-05 |
| f__Muribaculaceae | 1.74 | 0.445 | 1.03E-04 | 4.73E-04 |
| f__Acholeplasmataceae | -4.11 | 1.35 | 0.00243 | 0.00769 |
| f__Akkermansiaceae | 3.25 | 1.1 | 0.00341 | 0.0104 |
| g__Muribaculum | -6.11 | 0.798 | 8.55E-14 | 1.81E-12 |
| g__ASF356 | -4.1 | 0.815 | 6.61E-07 | 5.29E-06 |
| g__Lactobacillus | -4.28 | 0.855 | 7.64E-07 | 6.05E-06 |
| g__Ileibacterium | -4.2 | 0.862 | 1.42E-06 | 1.02E-05 |
| g__Eubacterium_coprostanoligenes_group | 3.16 | 0.67 | 3.17E-06 | 2.11E-05 |
| g__Dorea | -3.11 | 0.684 | 6.72E-06 | 4.21E-05 |
| g__Lachnospiraceae_FCS020_group | -3.82 | 0.853 | 9.05E-06 | 5.46E-05 |

| | | | | |
|---|---|---|---|---|
| g__Mucispirillum | 4.75 | 1.07 | 1.19E-05 | 6.94E-05 |
| g__Lachnospiraceae_UCG_006 | -3.34 | 0.792 | 2.87E-05 | 1.49E-04 |
| g__Muribaculaceae | 1.79 | 0.445 | 6.24E-05 | 2.92E-04 |
| g__Anaeroplasma | -4.11 | 1.35 | 0.00243 | 0.00756 |
| g__Roseburia | -2.8 | 0.931 | 0.00279 | 0.0086 |
| g__Akkermansia | 3.25 | 1.1 | 0.00341 | 0.0103 |
| g__Eubacterium_xylanophilum_group | -1.81 | 0.697 | 0.00984 | 0.0257 |
| g__Intestinimonas | -2.09 | 0.844 | 0.0134 | 0.0343 |
| uncultured_Bacteroidales | 2.26 | 0.459 | 1.19E-06 | 7.39E-06 |
| Ileibacterium_valens | -4.2 | 0.862 | 1.42E-06 | 8.49E-06 |
| mouse_gut | -2.9 | 0.747 | 1.19E-04 | 4.63E-04 |

**Legend:** d (domain), p (phylum), c (class), o (order), f (family), g (genus), standard error (St. Error), false-discovery rate adjusted p-value (FDR)

**Table S7**. Differences in gut microbiome taxa relative to young mice (showing taxa with q<0.05).

| Name | Log2FC | St.Error | P-value | FDR |
|---|---|---|---|---|
| Young/Aged | | | | |
| p__Firmicutes | -0.992 | 0.24 | 4.18E-05 | 1.70E-04 |
| p__Deferribacterota | 2.3 | 0.567 | 5.63E-05 | 2.22E-04 |
| p__Actinobacteriota | 1.6 | 0.396 | 6.25E-05 | 2.39E-04 |
| c__Bacilli | -2.51 | 0.506 | 9.01E-07 | 7.30E-06 |
| c__Deferribacteres | 2.3 | 0.567 | 5.63E-05 | 2.53E-04 |
| c__Coriobacteriia | 1.6 | 0.396 | 6.25E-05 | 2.74E-04 |
| c__Gammaproteobacteria | 1.62 | 0.409 | 8.68E-05 | 3.70E-04 |
| o__Acholeplasmatales | -4.48 | 0.712 | 6.63E-10 | 9.42E-09 |
| o__Lactobacillales | 2.38 | 0.445 | 1.24E-07 | 1.08E-06 |
| o__Clostridia_UCG_014 | -1.89 | 0.467 | 5.74E-05 | 2.92E-04 |
| o__Deferribacterales | 2.3 | 0.567 | 5.63E-05 | 2.92E-04 |
| o__Coriobacteriales | 1.6 | 0.396 | 6.25E-05 | 3.12E-04 |
| o__Burkholderiales | 1.62 | 0.409 | 8.68E-05 | 4.18E-04 |
| o__Clostridia_vadinBB60_group | -1.35 | 0.406 | 9.02E-04 | 0.00317 |
| o__Erysipelotrichales | 1.31 | 0.441 | 0.0032 | 0.0096 |
| f__Eubacterium_coprostanoligenes_group | 2.54 | 0.354 | 2.34E-12 | 3.57E-11 |
| f__Acholeplasmataceae | -4.48 | 0.712 | 6.63E-10 | 8.75E-09 |
| f__Rikenellaceae | -2.54 | 0.426 | 4.58E-09 | 5.18E-08 |
| f__Lactobacillaceae | 2.45 | 0.452 | 8.40E-08 | 7.92E-07 |
| f__Deferribacteraceae | 2.3 | 0.567 | 5.63E-05 | 2.93E-04 |
| f__Clostridia_UCG_014 | -1.89 | 0.467 | 5.74E-05 | 2.95E-04 |
| f__Atopobiaceae | 1.6 | 0.396 | 6.25E-05 | 3.13E-04 |
| f__Sutterellaceae | 1.62 | 0.409 | 8.68E-05 | 4.19E-04 |
| f__Clostridia_vadinBB60_group | -1.35 | 0.406 | 9.02E-04 | 0.00322 |
| f__Erysipelotrichaceae | 1.31 | 0.441 | 0.0032 | 0.00982 |

| | | | | |
|---|---|---|---|---|
| f__Bacteroidaceae | 0.698 | 0.246 | 0.00481 | 0.0141 |
| g__Muribaculum | -4.93 | 0.422 | 2.32E-28 | 1.19E-26 |
| g__Eubacterium_coprostanoligenes_group | 2.54 | 0.354 | 2.34E-12 | 4.02E-11 |
| g__Desulfovibrio | 2.59 | 0.39 | 7.79E-11 | 1.15E-09 |
| g__Anaeroplasma | -4.48 | 0.712 | 6.63E-10 | 8.84E-09 |
| g__Alistipes | -2.54 | 0.426 | 4.58E-09 | 5.78E-08 |
| g__Lactobacillus | 2.45 | 0.452 | 8.40E-08 | 8.40E-07 |
| g__ASF356 | -2.3 | 0.43 | 1.32E-07 | 1.25E-06 |
| g__Mucispirillum | 2.3 | 0.567 | 5.63E-05 | 2.72E-04 |
| g__Clostridia_UCG_014 | -1.89 | 0.467 | 5.74E-05 | 2.75E-04 |
| g__Coriobacteriaceae_UCG_002 | 1.6 | 0.396 | 6.25E-05 | 2.96E-04 |
| g__Parasutterella | 1.62 | 0.409 | 8.68E-05 | 3.95E-04 |
| g__Lachnoclostridium | 1.37 | 0.389 | 4.68E-04 | 0.00175 |
| g__Lachnospiraceae_FCS020_group | 1.55 | 0.45 | 6.08E-04 | 0.00222 |
| g__Clostridia_vadinBB60_group | -1.35 | 0.406 | 9.02E-04 | 0.00308 |
| g__Oscillibacter | -1.32 | 0.399 | 9.47E-04 | 0.0032 |
| g__Eubacterium_xylanophilum_group | 1.2 | 0.368 | 0.00116 | 0.00384 |
| g__GCA_900066575 | -1.25 | 0.42 | 0.00319 | 0.00961 |
| g__Bacteroides | 0.698 | 0.246 | 0.00481 | 0.0138 |
| g__Lachnospiraceae_UCG_001 | -1.63 | 0.586 | 0.00571 | 0.0159 |
| Not_Assigned | 0.673 | 0.263 | 0.0108 | 0.0273 |
| g__Faecalibaculum | 1.07 | 0.426 | 0.0126 | 0.0318 |
| uncultured_Barnesiella | 2.98 | 0.393 | 1.56E-13 | 2.44E-12 |
| uncultured_Bacteroidales | 1.53 | 0.243 | 5.94E-10 | 5.88E-09 |
| uncultured_bacterium | -0.66 | 0.139 | 2.52E-06 | 1.42E-05 |
| Not_Assigned | 0.432 | 0.103 | 2.93E-05 | 1.35E-04 |
| unidentified | -1.37 | 0.369 | 2.28E-04 | 8.52E-04 |
| mouse_gut | 0.998 | 0.395 | 0.0117 | 0.0264 |

**Legend:** d (domain), p (phylum), c (class), o (order), f (family), g (genus), standard error (St. Error), false-discovery rate adjusted p-value (FDR)

**Table S8**. Differences in gut microbiome taxa between sexes relative to male mice (showing taxa with q<0.05).

| Name | Log2FC | St.Error | P-value | FDR |
|---|---|---|---|---|
| Male/female | | | | |
| p__Deferribacterota | 2.03 | 0.479 | 2.52E-05 | 1.09E-04 |
| p__Desulfobacterota | -0.889 | 0.262 | 7.39E-04 | 0.00227 |
| p__Actinobacteriota | -1.12 | 0.334 | 8.55E-04 | 0.00256 |
| c__Deferribacteres | 2.03 | 0.479 | 2.52E-05 | 1.24E-04 |
| c__Desulfovibrionia | -0.889 | 0.262 | 7.39E-04 | 0.00255 |
| c__Coriobacteriia | -1.12 | 0.334 | 8.55E-04 | 0.00283 |
| c__Alphaproteobacteria | 1.23 | 0.472 | 0.00966 | 0.0257 |
| o__Clostridia_vadinBB60_group | 1.85 | 0.342 | 9.84E-08 | 8.86E-07 |
| o__Deferribacterales | 2.03 | 0.479 | 2.52E-05 | 1.39E-04 |
| o__Desulfovibrionales | -0.889 | 0.262 | 7.39E-04 | 0.00273 |
| o__Coriobacteriales | -1.12 | 0.334 | 8.55E-04 | 0.00308 |
| o__Rhodospirillales | 1.23 | 0.472 | 0.00966 | 0.0269 |
| f__Clostridia_vadinBB60_group | 1.85 | 0.342 | 9.84E-08 | 8.86E-07 |
| f__Deferribacteraceae | 2.03 | 0.479 | 2.52E-05 | 1.41E-04 |
| f__Desulfovibrionaceae | -0.889 | 0.262 | 7.39E-04 | 0.00276 |
| f__Atopobiaceae | -1.12 | 0.334 | 8.55E-04 | 0.00309 |
| f__Eubacterium_coprostanoligenes_group | -0.849 | 0.299 | 0.00464 | 0.0137 |
| f__uncultured | 1.23 | 0.472 | 0.00966 | 0.0268 |
| f__Rikenellaceae | 0.883 | 0.36 | 0.0144 | 0.037 |
| g__Roseburia | -2.3 | 0.415 | 4.70E-08 | 4.98E-07 |
| g__Clostridia_vadinBB60_group | 1.85 | 0.342 | 9.84E-08 | 9.58E-07 |
| g__Intestinimonas | -1.95 | 0.376 | 3.18E-07 | 2.63E-06 |
| g__Mucispirillum | 2.03 | 0.479 | 2.52E-05 | 1.33E-04 |
| g__Lachnospiraceae_UCG_001 | -1.97 | 0.494 | 7.53E-05 | 3.52E-04 |
| g__Anaerotruncus | -1.49 | 0.39 | 1.48E-04 | 6.38E-04 |
| g__Bilophila | -1.07 | 0.318 | 8.49E-04 | 0.00297 |

| | | | | |
|---|---|---|---|---|
| g__Coriobacteriaceae_UCG_002 | -1.12 | 0.334 | 8.55E-04 | 0.00297 |
| g__Incertae_Sedis | -0.961 | 0.296 | 0.00121 | 0.00399 |
| g__Lachnoclostridium | -1.03 | 0.328 | 0.00186 | 0.00581 |
| g__Eubacterium_coprostanoligenes_group | -0.849 | 0.299 | 0.00464 | 0.0135 |
| g__Blautia | -1.06 | 0.406 | 0.0093 | 0.0239 |
| g__Eubacterium_xylanophilum_group | -0.788 | 0.311 | 0.0115 | 0.029 |
| g__Alistipes | 0.883 | 0.36 | 0.0144 | 0.0356 |
| Clostridium_leptum | 1.93 | 0.34 | 2.32E-08 | 1.79E-07 |
| Lachnospiraceae_bacterium | -1.06 | 0.406 | 0.0093 | 0.0222 |

**Legend:** d (domain), p (phylum), c (class), o (order), f (family), g (genus), standard error (St. Error), false-discovery rate adjusted p-value (FDR)

**Table S9**. Pairwise comparisons between the diets for Bray-Curtis index (q<0.05).

| Comparisons | F-value | R-squared | P-value | FDR |
|---|---|---|---|---|
| Normal_chow vs High_fibre | 108.68 | 0.19315 | 0.001 | 0.00125 |
| Normal_chow vs No_fibre | 108.74 | 0.18886 | 0.001 | 0.00125 |
| Normal_chow vs AING93 | 46.572 | 0.097315 | 0.001 | 0.00125 |
| Normal_chow vs HAMSAB | 7.0146 | 0.016984 | 0.001 | 0.00125 |
| High_fibre vs No_fibre | 26.283 | 0.17606 | 0.001 | 0.00125 |
| High_fibre vs AING93 | 13.707 | 0.13477 | 0.001 | 0.00125 |
| High_fibre vs HAMSAB | 4.117 | 0.062269 | 0.001 | 0.00125 |
| No_fibre vs AING93 | 23.149 | 0.18646 | 0.001 | 0.00125 |
| No_fibre vs HAMSAB | 4.496 | 0.056557 | 0.002 | 0.002222 |
| AING93 vs HAMSAB | 2.3348 | 0.055151 | 0.044 | 0.044 |

**Table S10**. Differences in gut microbiome taxa across diets relative to the control (normal chow) diet (showing taxa with q<0.05).

| Taxa | Log2FC | St.Error | P-value | FDR |
|---|---|---|---|---|
| Normal chow/AIN93G | | | | |
| p__Deferribacterota | 3.47 | 0.752 | 4.93E-06 | 2.82E-05 |
| p__Desulfobacterota | -1.53 | 0.411 | 2.30E-04 | 8.06E-04 |
| c__Deferribacteres | 3.47 | 0.752 | 4.93E-06 | 3.07E-05 |
| c__Gammaproteobacteria | 2.12 | 0.542 | 1.02E-04 | 4.15E-04 |
| c__Desulfovibrionia | -1.53 | 0.411 | 2.30E-04 | 8.88E-04 |
| o__Deferribacterales | 3.47 | 0.752 | 4.93E-06 | 3.25E-05 |
| o__Burkholderiales | 2.12 | 0.542 | 1.02E-04 | 4.77E-04 |
| o__Desulfovibrionales | -1.53 | 0.411 | 2.30E-04 | 9.71E-04 |
| o__Erysipelotrichales | -2.16 | 0.585 | 2.51E-04 | 1.04E-03 |
| o__Clostridia_UCG_014 | 1.5 | 0.618 | 1.56E-02 | 3.93E-02 |
| f__Streptococcaceae | -2.36 | 0.406 | 1.02E-08 | 1.08E-07 |
| f__Eubacterium_coprostanoligenes_group | 2.45 | 0.469 | 2.65E-07 | 2.18E-06 |
| f__Deferribacteraceae | 3.47 | 0.752 | 4.93E-06 | 3.37E-05 |
| f__Butyricicoccaceae | 1.93 | 0.453 | 2.44E-05 | 1.38E-04 |
| f__Muribaculaceae | 1.3 | 0.311 | 3.66E-05 | 1.99E-04 |
| f__Bacteroidaceae | -1.3 | 0.327 | 8.27E-05 | 4.04E-04 |
| f__Sutterellaceae | 2.12 | 0.542 | 1.02E-04 | 4.83E-04 |
| f__Desulfovibrionaceae | -1.53 | 0.411 | 2.30E-04 | 9.80E-04 |
| f__Erysipelotrichaceae | -2.16 | 0.585 | 2.51E-04 | 1.06E-03 |
| f__Tannerellaceae | -1.55 | 0.447 | 5.55E-04 | 2.17E-03 |
| f__Clostridia_UCG_014 | 1.5 | 0.618 | 1.56E-02 | 3.90E-02 |
| g__Lactococcus | -2.36 | 0.406 | 1.02E-08 | 1.22E-07 |
| g__Bilophila | -2.88 | 0.499 | 1.29E-08 | 1.50E-07 |
| g__Roseburia | 3.52 | 0.652 | 9.49E-08 | 9.36E-07 |
| g__Dorea | 2.54 | 0.478 | 1.58E-07 | 1.46E-06 |

| | | | | |
|---|---|---|---|---|
| g__Eubacterium_coprostanoligenes_group | 2.45 | 0.469 | 2.65E-07 | 2.24E-06 |
| g__Lachnospiraceae_UCG_006 | 2.81 | 0.554 | 5.62E-07 | 4.55E-06 |
| g__Eubacterium_xylanophilum_group | 2.31 | 0.488 | 2.89E-06 | 1.95E-05 |
| g__Mucispirillum | 3.47 | 0.752 | 4.93E-06 | 3.29E-05 |
| g__Tuzzerella | -2.21 | 0.505 | 1.42E-05 | 8.24E-05 |
| g__UCG_009 | 1.93 | 0.453 | 2.44E-05 | 1.30E-04 |
| g__Muribaculaceae | 1.31 | 0.311 | 2.91E-05 | 1.52E-04 |
| g__Faecalibaculum | -2.35 | 0.564 | 3.74E-05 | 1.92E-04 |
| g__Bacteroides | -1.3 | 0.327 | 8.27E-05 | 3.81E-04 |
| g__Parasutterella | 2.12 | 0.542 | 1.02E-04 | 4.61E-04 |
| g__Lachnospiraceae_FCS020_group | 2.18 | 0.597 | 2.84E-04 | 1.15E-03 |
| g__Lachnospiraceae_UCG_001 | 2.2 | 0.776 | 4.73E-03 | 1.36E-02 |
| g__uncultured | -1.11 | 0.416 | 8.00E-03 | 0.021 |
| g__Blautia | -1.69 | 0.638 | 0.00845 | 0.0221 |
| g__Clostridia_UCG_014 | 1.5 | 0.618 | 0.0156 | 0.0375 |
| Clostridium_leptum | 3.44 | 0.533 | 2.38E-10 | 2.62E-09 |
| Lactococcus_lactis | -2.36 | 0.406 | 1.02E-08 | 8.45E-08 |
| uncultured_Bacteroidales | 1.65 | 0.321 | 3.85E-07 | 2.38E-06 |
| uncultured_Barnesiella | 2.29 | 0.521 | 1.36E-05 | 6.73E-05 |
| Lachnospiraceae_bacterium | -1.69 | 0.638 | 8.45E-03 | 2.04E-02 |
| Normal chow/HAMSAB | | | | |
| o__Peptococcales | 2.14 | 0.71 | 2.73E-03 | 8.66E-03 |
| f__Peptococcaceae | 2.14 | 0.71 | 2.73E-03 | 9.55E-03 |
| f__Muribaculaceae | 1.19 | 0.5 | 1.73E-02 | 4.54E-02 |
| g__Bilophila | -2.5 | 0.801 | 1.89E-03 | 6.32E-03 |
| g__Blautia | -2.96 | 1.02 | 4.07E-03 | 1.28E-02 |
| g__Anaerotruncus | -2.52 | 0.984 | 1.07E-02 | 2.89E-02 |
| g__GCA_900066575 | -2.27 | 0.894 | 1.15E-02 | 3.06E-02 |
| g__Muribaculaceae | 1.23 | 0.5 | 1.45E-02 | 3.75E-02 |
| g__Dorea | 1.82 | 0.768 | 1.83E-02 | 4.51E-02 |

| | | | | |
|---|---|---|---|---|
| uncultured_Barnesiella | 3.77 | 0.836 | 7.80E-06 | 4.54E-05 |
| uncultured_Bacteroidales | 2.06 | 0.516 | 7.68E-05 | 3.54E-04 |
| Lachnospiraceae_bacterium | -2.96 | 1.02 | 4.07E-03 | 0.0129 |
| Normal chow/High-fibre | | | | |
| p__Verrucomicrobiota | -5.65 | 0.83 | 2.72E-11 | 3.11E-10 |
| p__Deferribacterota | 3.7 | 0.808 | 5.62E-06 | 2.90E-05 |
| p__Bacteroidota | -0.567 | 0.145 | 1.03E-04 | 3.61E-04 |
| p__Proteobacteria | -1.53 | 0.588 | 9.29E-03 | 2.30E-02 |
| c__Verrucomicrobiae | -5.65 | 0.83 | 2.72E-11 | 3.67E-10 |
| c__Deferribacteres | 3.7 | 0.808 | 5.62E-06 | 3.21E-05 |
| c__Bacteroidia | -0.567 | 0.145 | 1.03E-04 | 4.08E-04 |
| o__Verrucomicrobiales | -5.65 | 0.83 | 2.72E-11 | 4.08E-10 |
| o__Acholeplasmatales | 4.68 | 1.01 | 4.84E-06 | 3.18E-05 |
| o__Deferribacterales | 3.7 | 0.808 | 5.62E-06 | 3.53E-05 |
| o__Bacteroidales | -0.567 | 0.145 | 1.03E-04 | 4.57E-04 |
| o__Clostridia_vadinBB60_group | 1.82 | 0.578 | 0.00174 | 0.00546 |
| o__Peptococcales | -1.49 | 0.475 | 0.00177 | 0.00549 |
| o__Oscillospirales | 1.12 | 0.484 | 0.0213 | 0.0472 |
| f__Bacteroidaceae | -3.08 | 0.351 | 2.30E-17 | 5.35E-16 |
| f__Butyricicoccaceae | 3.71 | 0.487 | 1.14E-13 | 1.96E-12 |
| f__Akkermansiaceae | -5.65 | 0.83 | 2.72E-11 | 3.36E-10 |
| f__Eubacterium_coprostanoligenes_group | 2.96 | 0.504 | 7.19E-09 | 6.78E-08 |
| f__Muribaculaceae | 1.89 | 0.335 | 2.62E-08 | 2.20E-07 |
| f__Acholeplasmataceae | 4.68 | 1.01 | 4.84E-06 | 3.04E-05 |
| f__Deferribacteraceae | 3.7 | 0.808 | 5.62E-06 | 3.35E-05 |
| f__Tannerellaceae | -2.03 | 0.481 | 2.81E-05 | 1.39E-04 |
| f__Lactobacillaceae | 2.37 | 0.643 | 2.47E-04 | 9.99E-04 |
| f__Rikenellaceae | 2.11 | 0.607 | 5.66E-04 | 0.00215 |
| f__Clostridia_vadinBB60_group | 1.82 | 0.578 | 0.00174 | 0.00547 |
| f__Peptococcaceae | -1.49 | 0.475 | 0.00177 | 0.00552 |

| | | | | |
|---|---|---|---|---|
| f__Oscillospiraceae | 1.16 | 0.487 | 0.0171 | 0.0394 |
| g__Bacteroides | -3.08 | 0.351 | 2.30E-17 | 6.12E-16 |
| g__Roseburia | 5.47 | 0.7 | 2.94E-14 | 6.22E-13 |
| g__UCG_009 | 3.71 | 0.487 | 1.14E-13 | 2.11E-12 |
| g__Lachnospiraceae_UCG_006 | 4.33 | 0.596 | 1.27E-12 | 1.95E-11 |
| g__A2 | 4.83 | 0.683 | 4.66E-12 | 6.72E-11 |
| g__Eubacterium_xylanophilum_group | 3.67 | 0.524 | 7.38E-12 | 1.02E-10 |
| g__Akkermansia | -5.65 | 0.83 | 2.72E-11 | 3.48E-10 |
| g__Dorea | 3.47 | 0.514 | 3.66E-11 | 4.48E-10 |
| g__Eubacterium_coprostanoligenes_group | 2.96 | 0.504 | 7.19E-09 | 7.19E-08 |
| g__Muribaculaceae | 1.88 | 0.334 | 2.99E-08 | 2.76E-07 |
| g__Desulfovibrio | 3.03 | 0.555 | 7.06E-08 | 6.05E-07 |
| g__Lachnospiraceae_FCS020_group | 3.24 | 0.641 | 6.11E-07 | 4.35E-06 |
| g__Anaeroplasma | 4.68 | 1.01 | 4.84E-06 | 2.93E-05 |
| g__Mucispirillum | 3.7 | 0.808 | 5.62E-06 | 3.32E-05 |
| g__Tuzzerella | 2.46 | 0.543 | 6.83E-06 | 3.90E-05 |
| g__Lachnospiraceae_UCG_001 | 3.57 | 0.834 | 2.15E-05 | 1.07E-04 |
| g__Intestinimonas | 2.6 | 0.635 | 4.85E-05 | 2.24E-04 |
| g__Lactobacillus | 2.37 | 0.643 | 2.47E-04 | 9.73E-04 |
| g__Oscillibacter | 1.97 | 0.567 | 5.46E-04 | 0.0019 |
| g__Alistipes | 2.11 | 0.607 | 5.66E-04 | 0.00195 |
| g__Faecalibaculum | -1.95 | 0.606 | 0.00138 | 0.00408 |
| g__Lachnoclostridium | 1.75 | 0.554 | 0.0017 | 0.00494 |
| g__Clostridia_vadinBB60_group | 1.82 | 0.578 | 0.00174 | 0.00503 |
| g__Incertae_Sedis | 1.42 | 0.499 | 0.00448 | 0.012 |
| g__Dubosiella | 0.994 | 0.425 | 0.0197 | 0.0424 |
| uncultured_Bacteroidales | 4.57 | 0.345 | 6.36E-35 | 3.15E-33 |
| Clostridium_leptum | 5.66 | 0.573 | 2.73E-21 | 6.01E-20 |
| Not_Assigned | -0.737 | 0.146 | 6.22E-07 | 3.62E-06 |
| unidentified | 2.17 | 0.526 | 4.35E-05 | 1.83E-04 |

| | | | | |
|---|---|---|---|---|
| mouse_gut | 1.77 | 0.562 | 0.0017 | 0.00475 |
| uncultured_bacterium | 0.594 | 0.197 | 0.00275 | 0.00736 |
| uncultured_Barnesiella | 1.67 | 0.56 | 0.00292 | 0.00771 |
| Dubosiella_newyorkensis | 0.994 | 0.425 | 0.0197 | 0.0395 |
| Normal chow/low-fibre | | | | |
| p__Desulfobacterota | -3.93 | 0.455 | 5.79E-17 | 1.22E-15 |
| p__Deferribacterota | 5.74 | 0.831 | 1.40E-11 | 1.60E-10 |
| p__Verrucomicrobiota | -5.63 | 0.854 | 1.02E-10 | 1.07E-09 |
| p__Actinobacteriota | -2.53 | 0.58 | 1.53E-05 | 6.43E-05 |
| p__Bacteroidota | -0.519 | 0.149 | 5.42E-04 | 0.00159 |
| p__Firmicutes | 1.04 | 0.352 | 0.00323 | 0.00754 |
| c__Desulfovibrionia | -3.93 | 0.455 | 5.79E-17 | 1.56E-15 |
| c__Deferribacteres | 5.74 | 0.831 | 1.40E-11 | 1.89E-10 |
| c__Verrucomicrobiae | -5.63 | 0.854 | 1.02E-10 | 1.27E-09 |
| c__Coriobacteriia | -2.53 | 0.58 | 1.53E-05 | 7.30E-05 |
| c__Clostridia | 1.52 | 0.423 | 3.57E-04 | 0.0012 |
| c__Bacteroidia | -0.519 | 0.149 | 5.42E-04 | 0.00169 |
| c__Gammaproteobacteria | 1.74 | 0.599 | 0.0039 | 0.00987 |
| o__Desulfovibrionales | -3.93 | 0.455 | 5.79E-17 | 1.56E-15 |
| o__Deferribacterales | 5.74 | 0.831 | 1.40E-11 | 2.10E-10 |
| o__Verrucomicrobiales | -5.63 | 0.854 | 1.02E-10 | 1.37E-09 |
| o__Lachnospirales | 2.43 | 0.477 | 4.86E-07 | 3.36E-06 |
| o__Clostridia_vadinBB60_group | 2.98 | 0.594 | 7.35E-07 | 4.84E-06 |
| o__Coriobacteriales | -2.53 | 0.58 | 1.53E-05 | 7.95E-05 |
| o__Clostridia_UCG_014 | 2.8 | 0.683 | 4.94E-05 | 2.30E-04 |
| o__Bacteroidales | -0.519 | 0.149 | 5.42E-04 | 0.00181 |
| o__Lactobacillales | -2.26 | 0.652 | 5.52E-04 | 0.00182 |
| o__Erysipelotrichales | -2.12 | 0.647 | 0.00112 | 0.00338 |
| o__Burkholderiales | 1.74 | 0.599 | 0.0039 | 0.0101 |
| o__Acholeplasmatales | 2.77 | 1.04 | 0.00828 | 0.0205 |

| | | | | |
|---|---|---|---|---|
| o__Peptococcales | -1.14 | 0.489 | 0.0206 | 0.0438 |
| f__Tannerellaceae | -4.46 | 0.494 | 3.00E-18 | 7.43E-17 |
| f__Desulfovibrionaceae | -3.93 | 0.455 | 5.79E-17 | 1.27E-15 |
| f__Bacteroidaceae | -2.55 | 0.361 | 4.89E-12 | 6.24E-11 |
| f__Deferribacteraceae | 5.74 | 0.831 | 1.40E-11 | 1.73E-10 |
| f__Akkermansiaceae | -5.63 | 0.854 | 1.02E-10 | 1.22E-09 |
| f__Streptococcaceae | -2.88 | 0.449 | 2.76E-10 | 3.22E-09 |
| f__Lachnospiraceae | 2.43 | 0.477 | 4.86E-07 | 3.38E-06 |
| f__Clostridia_vadinBB60_group | 2.98 | 0.594 | 7.35E-07 | 4.93E-06 |
| f__Eubacterium_coprostanoligenes_group | 2.58 | 0.519 | 8.90E-07 | 5.88E-06 |
| f__Atopobiaceae | -2.53 | 0.58 | 1.53E-05 | 7.98E-05 |
| f__Clostridia_UCG_014 | 2.8 | 0.683 | 4.94E-05 | 2.35E-04 |
| f__Erysipelotrichaceae | -2.12 | 0.647 | 0.00112 | 0.00371 |
| f__Muribaculaceae | 1.12 | 0.344 | 0.00127 | 0.00399 |
| f__Sutterellaceae | 1.74 | 0.599 | 0.0039 | 0.011 |
| f__Acholeplasmataceae | 2.77 | 1.04 | 0.00828 | 0.0216 |
| f__Peptococcaceae | -1.14 | 0.489 | 0.0206 | 0.0464 |
| g__Bilophila | -5.95 | 0.552 | 9.87E-25 | 3.74E-23 |
| g__Roseburia | 6.13 | 0.72 | 1.65E-16 | 4.10E-15 |
| g__Lachnospiraceae_NK4A136_group | 4.89 | 0.58 | 2.83E-16 | 6.57E-15 |
| g__Lachnospiraceae_FCS020_group | 5.34 | 0.66 | 3.60E-15 | 7.63E-14 |
| g__Bacteroides | -2.55 | 0.361 | 4.89E-12 | 7.33E-11 |
| g__Mucispirillum | 5.74 | 0.831 | 1.40E-11 | 2.01E-10 |
| g__A2 | 4.67 | 0.703 | 7.41E-11 | 9.70E-10 |
| g__Akkermansia | -5.63 | 0.854 | 1.02E-10 | 1.26E-09 |
| g__Faecalibaculum | -4.03 | 0.624 | 2.29E-10 | 2.80E-09 |
| g__Lactococcus | -2.88 | 0.449 | 2.76E-10 | 3.30E-09 |
| g__Lachnospiraceae_UCG_001 | 4.91 | 0.858 | 1.75E-08 | 1.64E-07 |
| g__Desulfovibrio | 2.93 | 0.571 | 3.94E-07 | 2.90E-06 |
| g__Dorea | 2.67 | 0.529 | 5.87E-07 | 4.22E-06 |

| | | | | |
|---|---|---|---|---|
| g__Clostridia_vadinBB60_group | 2.98 | 0.594 | 7.35E-07 | 5.14E-06 |
| g__Eubacterium_coprostanoligenes_group | 2.58 | 0.519 | 8.90E-07 | 6.05E-06 |
| g__Coriobacteriaceae_UCG_002 | -2.53 | 0.58 | 1.53E-05 | 7.96E-05 |
| g__Dubosiella | -1.91 | 0.437 | 1.54E-05 | 7.96E-05 |
| g__Clostridia_UCG_014 | 2.8 | 0.683 | 4.94E-05 | 2.26E-04 |
| g__Eubacterium_xylanophilum_group | 2.16 | 0.539 | 6.91E-05 | 3.02E-04 |
| g__Tuzzerella | -1.86 | 0.558 | 8.98E-04 | 2.92E-03 |
| g__Muribaculaceae | 1.1 | 0.344 | 1.43E-03 | 4.35E-03 |
| g__Parasutterella | 1.74 | 0.599 | 3.90E-03 | 1.08E-02 |
| g__Anaeroplasma | 2.77 | 1.04 | 8.28E-03 | 2.08E-02 |
| g__ASF356 | 1.65 | 0.63 | 9.06E-03 | 2.25E-02 |
| Not_Assigned | -0.88 | 0.385 | 2.29E-02 | 4.97E-02 |
| Clostridium_leptum | 5.58 | 0.59 | 9.26E-20 | 2.29E-18 |
| uncultured_Barnesiella | 5.08 | 0.576 | 1.49E-17 | 3.22E-16 |
| mouse_gut | 4.03 | 0.578 | 9.61E-12 | 9.52E-11 |
| Lactococcus_lactis | -2.88 | 0.449 | 2.76E-10 | 2.49E-09 |
| uncultured_Bacteroidales | 1.56 | 0.355 | 1.35E-05 | 6.23E-05 |
| Dubosiella_newyorkensis | -1.91 | 0.437 | 1.54E-05 | 6.76E-05 |
| Not_Assigned | -0.588 | 0.15 | 1.02E-04 | 3.66E-04 |
| unidentified | 2.02 | 0.541 | 2.03E-04 | 6.80E-04 |

**Legend:** d (domain), p (phylum), c (class), o (order), f (family), g (genus), standard error (St. Error), false-discovery rate adjusted p-value (FDR)

**Table S11**. Differences in gut microbiome taxa between the compartments relative to the large intestine (showing taxa with q<0.05).

| Name | Log2FC | St.Error | P-value | FDR |
|---|---|---|---|---|
| Large intestine/small intestine | | | | |
| p__Firmicutes | 4.06 | 0.217 | 1.35E-60 | 1.70E-58 |
| p__Deferribacterota | 4.27 | 0.514 | 7.07E-16 | 1.48E-14 |
| p__Bacteroidota | -0.448 | 0.0922 | 1.57E-06 | 9.90E-06 |
| p__Proteobacteria | 1.56 | 0.374 | 3.66E-05 | 1.54E-04 |
| c__Clostridia | 5.27 | 0.262 | 4.91E-68 | 7.96E-66 |
| c__Deferribacteres | 4.27 | 0.514 | 7.07E-16 | 1.91E-14 |
| c__Alphaproteobacteria | 3.11 | 0.506 | 1.65E-09 | 2.23E-08 |
| c__Bacteroidia | -0.448 | 0.0922 | 1.57E-06 | 1.11E-05 |
| o__Clostridia_vadinBB60_group | 7.9 | 0.367 | 8.79E-75 | 2.37E-72 |
| o__Oscillospirales | 6.05 | 0.308 | 1.82E-65 | 2.46E-63 |
| o__Lachnospirales | 5.05 | 0.295 | 5.91E-53 | 5.32E-51 |
| o__Clostridia_UCG_014 | 5.33 | 0.422 | 3.02E-32 | 2.04E-30 |
| o__Deferribacterales | 4.27 | 0.514 | 7.07E-16 | 1.91E-14 |
| o__Peptococcales | 2.3 | 0.302 | 1.33E-13 | 3.00E-12 |
| o__Rhodospirillales | 3.11 | 0.506 | 1.65E-09 | 2.13E-08 |
| o__Bacteroidales | -0.448 | 0.0922 | 1.57E-06 | 1.09E-05 |
| o__Lactobacillales | -1.55 | 0.403 | 1.36E-04 | 5.92E-04 |
| o__Acholeplasmatales | 2.06 | 0.645 | 0.00148 | 0.00483 |
| f__Bacteroidaceae | 6.06 | 0.223 | 2.07E-103 | 8.19E-101 |
| f__Clostridia_vadinBB60_group | 7.9 | 0.367 | 8.79E-75 | 1.74E-72 |
| f__Oscillospiraceae | 5.49 | 0.31 | 5.83E-56 | 7.70E-54 |
| f__Lachnospiraceae | 5.05 | 0.295 | 5.91E-53 | 5.85E-51 |
| f__Tannerellaceae | 4.89 | 0.306 | 1.34E-47 | 1.06E-45 |
| f__Ruminococcaceae | 5.13 | 0.337 | 8.47E-44 | 5.59E-42 |
| f__Rikenellaceae | 5.29 | 0.386 | 5.97E-37 | 3.38E-35 |

| | | | | |
|---|---|---|---|---|
| f__Clostridia_UCG_014 | 5.33 | 0.422 | 3.02E-32 | 1.49E-30 |
| f__Eubacterium_coprostanoligenes_group | 3.6 | 0.321 | 1.88E-26 | 5.74E-25 |
| f__Deferribacteraceae | 4.27 | 0.514 | 7.07E-16 | 1.65E-14 |
| f__Butyricicoccaceae | 2.41 | 0.309 | 3.17E-14 | 6.76E-13 |
| f__Peptococcaceae | 2.3 | 0.302 | 1.33E-13 | 2.40E-12 |
| f__uncultured | 3.11 | 0.506 | 1.65E-09 | 2.05E-08 |
| f__Lactobacillaceae | -1.58 | 0.409 | 1.19E-04 | 5.50E-04 |
| f__Muribaculaceae | -0.736 | 0.213 | 5.85E-04 | 0.00227 |
| f__Acholeplasmataceae | 2.06 | 0.645 | 0.00148 | 0.0049 |
| g__Bacteroides | 6.06 | 0.223 | 2.07E-103 | 1.49E-100 |
| Not_Assigned | 6.08 | 0.238 | 2.33E-95 | 8.38E-93 |
| g__Clostridia_vadinBB60_group | 7.9 | 0.367 | 8.79E-75 | 2.11E-72 |
| g__uncultured | 5.04 | 0.285 | 7.02E-56 | 1.26E-53 |
| g__Alistipes | 5.29 | 0.386 | 5.97E-37 | 7.17E-35 |
| g__Oscillibacter | 4.69 | 0.361 | 7.92E-34 | 8.14E-32 |
| g__Clostridia_UCG_014 | 5.33 | 0.422 | 3.02E-32 | 2.72E-30 |
| g__Colidextribacter | 4.34 | 0.354 | 1.07E-30 | 8.59E-29 |
| g__Lachnospiraceae_NK4A136_group | 4.38 | 0.358 | 1.63E-30 | 1.17E-28 |
| g__Incertae_Sedis | 3.85 | 0.317 | 3.09E-30 | 2.03E-28 |
| g__Eubacterium_xylanophilum_group | 3.96 | 0.333 | 4.21E-29 | 2.53E-27 |
| g__Lachnoclostridium | 3.98 | 0.352 | 9.95E-27 | 4.22E-25 |
| g__Eubacterium_coprostanoligenes_group | 3.6 | 0.321 | 1.88E-26 | 7.53E-25 |
| g__Intestinimonas | 3.76 | 0.404 | 2.97E-19 | 9.71E-18 |
| g__Lachnospiraceae_FCS020_group | 3.51 | 0.408 | 8.09E-17 | 2.24E-15 |
| g__Bilophila | 2.89 | 0.341 | 1.98E-16 | 5.09E-15 |
| g__Mucispirillum | 4.27 | 0.514 | 7.07E-16 | 1.70E-14 |
| g__A2 | 3.42 | 0.434 | 1.97E-14 | 4.57E-13 |
| g__UCG_009 | 2.41 | 0.309 | 3.17E-14 | 7.08E-13 |
| g__GCA_900066575 | 2.67 | 0.381 | 7.12E-12 | 1.19E-10 |
| g__Lachnospiraceae_UCG_006 | 2.6 | 0.379 | 1.87E-11 | 3.07E-10 |

| | | | | |
|---|---|---|---|---|
| g__Roseburia | 2.83 | 0.445 | 4.14E-10 | 5.62E-09 |
| g__Dorea | 1.61 | 0.327 | 1.16E-06 | 8.60E-06 |
| g__Anaerotruncus | 1.85 | 0.419 | 1.27E-05 | 7.58E-05 |
| g__Lachnospiraceae_UCG_001 | 2.06 | 0.53 | 1.14E-04 | 5.05E-04 |
| g__Lactobacillus | -1.58 | 0.409 | 1.19E-04 | 5.24E-04 |
| g__Tuzzerella | 1.27 | 0.345 | 2.55E-04 | 0.00107 |
| g__Muribaculaceae | -0.74 | 0.213 | 5.46E-04 | 0.00201 |
| g__Anaeroplasma | 2.06 | 0.645 | 0.00148 | 0.00475 |
| g__Faecalibaculum | -1.07 | 0.386 | 0.00573 | 0.0159 |
| g__Blautia | 1.13 | 0.436 | 0.0101 | 0.0258 |
| unidentified | 5.05 | 0.334 | 2.90E-43 | 2.87E-41 |
| Not_Assigned | 0.931 | 0.0929 | 7.61E-22 | 2.15E-20 |
| mouse_gut | 2.7 | 0.357 | 1.92E-13 | 2.72E-12 |
| uncultured_Barnesiella | -1.73 | 0.356 | 1.46E-06 | 8.49E-06 |
| Clostridium_leptum | 1.34 | 0.364 | 2.72E-04 | 9.96E-04 |
| Lachnospiraceae_bacterium | 1.13 | 0.436 | 0.0101 | 0.0239 |
| uncultured_Bacteroidales | -0.561 | 0.22 | 0.011 | 0.0252 |

**Legend:** d (domain), p (phylum), c (class), o (order), f (family), g (genus), standard error (St. Error), false-discovery rate adjusted p-value (FDR)

**Table S12**. Pairwise comparisons between the sequencing batches for Bray-Curtis index.

| Pair | F-value | R-squared | P-value | FDR |
|---|---|---|---|---|
| May-21 vs Aug-21 | 8.6092 | 0.02644 | 0.001 | 0.001 |
| May-21 vs Nov-22 | 28.1154 | 0.202101 | 0.001 | 0.001 |
| May-21 vs Oct-20 | 13.1709 | 0.122896 | 0.001 | 0.001 |
| May-21 vs Dec-21 | 27.3616 | 0.200655 | 0.001 | 0.001 |
| May-21 vs Mar-20 | 36.8297 | 0.185233 | 0.001 | 0.001 |
| Aug-21 vs Nov-22 | 58.6442 | 0.158222 | 0.001 | 0.001 |
| Aug-21 vs Oct-20 | 27.5621 | 0.085447 | 0.001 | 0.001 |
| Aug-21 vs Dec-21 | 58.2847 | 0.15826 | 0.001 | 0.001 |
| Aug-21 vs Mar-20 | 15.4778 | 0.040895 | 0.001 | 0.001 |
| Nov-22 vs Oct-20 | 17.6923 | 0.165825 | 0.001 | 0.001 |
| Nov-22 vs Dec-21 | 23.9105 | 0.186932 | 0.001 | 0.001 |
| Nov-22 vs Mar-20 | 110.034 | 0.41206 | 0.001 | 0.001 |
| Oct-20 vs Dec-21 | 8.6798 | 0.090717 | 0.001 | 0.001 |
| Oct-20 vs Mar-20 | 56.8606 | 0.288837 | 0.001 | 0.001 |
| Dec-21 vs Mar-20 | 104.79 | 0.403363 | 0.001 | 0.001 |

**Table S13**. Differences in gut microbiome taxa between sequencing batches relative to Aug-21 batch (showing taxa with q<0.05).

| **Taxa** | **Log2FC** | **St. Error** | **P-value** | **FDR** |
|---|---|---|---|---|
| Aug21/Dec21 | | | | |
| p__Verrucomicrobiota | 7.83 | 0.876 | 6.06E-18 | 1.91E-16 |
| p__Deferribacterota | -6.14 | 0.852 | 1.99E-12 | 2.50E-11 |
| p__Bacteroidota | 0.833 | 0.153 | 7.68E-08 | 5.09E-07 |
| p__Actinobacteriota | -2.55 | 0.595 | 2.20E-05 | 8.40E-05 |
| p__Desulfobacterota | -1.83 | 0.466 | 9.45E-05 | 3.05E-04 |
| p__Firmicutes | -1.17 | 0.361 | 1.27E-03 | 0.00292 |
| c__Verrucomicrobiae | 7.83 | 0.876 | 6.06E-18 | 2.46E-16 |
| c__Deferribacteres | -6.14 | 0.852 | 1.99E-12 | 3.22E-11 |
| c__Bacteroidia | 0.833 | 0.153 | 7.68E-08 | 6.22E-07 |
| c__Coriobacteriia | -2.55 | 0.595 | 2.20E-05 | 9.63E-05 |
| c__Desulfovibrionia | -1.83 | 0.466 | 9.45E-05 | 3.64E-04 |
| c__Bacilli | -2.65 | 0.76 | 5.22E-04 | 0.00163 |
| c__Clostridia | -1.43 | 0.434 | 1.03E-03 | 2.82E-03 |
| c__Gammaproteobacteria | -1.48 | 0.614 | 1.62E-02 | 3.54E-02 |
| o__Verrucomicrobiales | 7.83 | 0.876 | 6.06E-18 | 2.34E-16 |
| o__Deferribacterales | -6.14 | 0.852 | 1.99E-12 | 3.83E-11 |
| o__Erysipelotrichales | -4.12 | 0.663 | 1.01E-09 | 1.43E-08 |
| o__Clostridia_UCG_014 | 3.97 | 0.701 | 2.36E-08 | 2.45E-07 |
| o__Bacteroidales | 0.833 | 0.153 | 7.68E-08 | 6.91E-07 |
| o__Coriobacteriales | -2.55 | 0.595 | 2.20E-05 | 1.10E-04 |
| o__Desulfovibrionales | -1.83 | 0.466 | 9.45E-05 | 4.11E-04 |
| o__Lachnospirales | -1.88 | 0.489 | 1.33E-04 | 5.48E-04 |
| o__Oscillospirales | -1.76 | 0.511 | 6.10E-04 | 2.11E-03 |
| o__Lactobacillales | -2.02 | 0.668 | 0.00267 | 0.00721 |
| o__Burkholderiales | -1.48 | 0.614 | 0.0162 | 0.0363 |

| | | | | |
|---|---|---|---|---|
| f__Akkermansiaceae | 7.83 | 0.876 | 6.06E-18 | 1.85E-16 |
| f__Eubacterium_coprostanoligenes_group | -4.07 | 0.532 | 8.87E-14 | 1.85E-12 |
| f__Deferribacteraceae | -6.14 | 0.852 | 1.99E-12 | 3.15E-11 |
| f__Streptococcaceae | -2.93 | 0.46 | 4.13E-10 | 5.28E-09 |
| f__Erysipelotrichaceae | -4.12 | 0.663 | 1.01E-09 | 1.21E-08 |
| f__Clostridia_UCG_014 | 3.97 | 0.701 | 2.36E-08 | 2.28E-07 |
| f__Atopobiaceae | -2.55 | 0.595 | 2.20E-05 | 1.15E-04 |
| f__Desulfovibrionaceae | -1.83 | 0.466 | 9.45E-05 | 4.25E-04 |
| f__Lachnospiraceae | -1.88 | 0.489 | 1.33E-04 | 5.73E-04 |
| f__Oscillospiraceae | -1.95 | 0.514 | 1.69E-04 | 6.90E-04 |
| f__Ruminococcaceae | -1.66 | 0.56 | 3.15E-03 | 9.16E-03 |
| f__Butyricicoccaceae | -1.32 | 0.513 | 1.01E-02 | 0.0261 |
| f__Sutterellaceae | -1.48 | 0.614 | 0.0162 | 0.039 |
| f__Muribaculaceae | 0.847 | 0.353 | 0.0167 | 0.0399 |
| g__Akkermansia | 7.83 | 0.876 | 6.06E-18 | 1.98E-16 |
| g__Blautia | -6.38 | 0.724 | 1.63E-17 | 5.09E-16 |
| g__Eubacterium_coprostanoligenes_group | -4.07 | 0.532 | 8.87E-14 | 1.93E-12 |
| g__Mucispirillum | -6.14 | 0.852 | 1.99E-12 | 3.67E-11 |
| g__Lactococcus | -2.93 | 0.46 | 4.13E-10 | 5.73E-09 |
| g__Clostridia_UCG_014 | 3.97 | 0.701 | 2.36E-08 | 2.70E-07 |
| g__Anaerotruncus | -3.84 | 0.695 | 5.04E-08 | 5.26E-07 |
| g__Desulfovibrio | -3.09 | 0.585 | 1.87E-07 | 1.68E-06 |
| g__Tuzzerella | -2.74 | 0.572 | 2.13E-06 | 1.48E-05 |
| g__Faecalibaculum | -2.86 | 0.639 | 9.10E-06 | 5.60E-05 |
| g__Coriobacteriaceae_UCG_002 | -2.55 | 0.595 | 2.20E-05 | 1.23E-04 |
| g__Colidextribacter | -2.44 | 0.588 | 3.80E-05 | 1.97E-04 |
| g__Lachnospiraceae_UCG_006 | -2.57 | 0.628 | 4.84E-05 | 2.45E-04 |
| g__Lachnospiraceae_FCS020_group | -2.72 | 0.677 | 6.61E-05 | 3.13E-04 |

| | | | | |
|---|---|---|---|---|
| g__ASF356 | -2.53 | 0.646 | 1.00E-04 | 4.55E-04 |
| g__Dubosiella | -1.74 | 0.448 | 1.12E-04 | 4.98E-04 |
| g__Bilophila | -2.13 | 0.565 | 1.87E-04 | 7.57E-04 |
| g__Ileibacterium | -2.49 | 0.684 | 2.89E-04 | 0.0011 |
| g__Muribaculum | -2.3 | 0.633 | 3.05E-04 | 0.00115 |
| g__Oscillibacter | -2.07 | 0.598 | 5.97E-04 | 0.00208 |
| g__uncultured | -1.62 | 0.472 | 6.63E-04 | 0.00226 |
| g__Lachnoclostridium | -1.97 | 0.584 | 8.11E-04 | 0.00269 |
| g__Roseburia | -2.38 | 0.739 | 0.00132 | 0.00407 |
| Not_Assigned | -1.21 | 0.395 | 0.00232 | 0.00686 |
| g__Incertae_Sedis | -1.53 | 0.526 | 0.00384 | 0.0109 |
| g__A2 | -1.98 | 0.72 | 0.00631 | 0.0162 |
| g__UCG_009 | -1.32 | 0.513 | 0.0101 | 0.0243 |
| g__Muribaculaceae | 0.86 | 0.353 | 0.0151 | 0.0344 |
| g__Parasutterella | -1.48 | 0.614 | 0.0162 | 0.0364 |
| Lachnospiraceae_bacterium | -6.38 | 0.724 | 1.63E-17 | 5.37E-16 |
| Lactococcus_lactis | -2.93 | 0.46 | 4.13E-10 | 5.12E-09 |
| Clostridium_leptum | -3.4 | 0.605 | 2.86E-08 | 2.36E-07 |
| mouse_gut | -3.05 | 0.593 | 3.73E-07 | 2.73E-06 |
| Dubosiella_newyorkensis | -1.74 | 0.448 | 1.12E-04 | 4.62E-04 |
| Ileibacterium_valens | -2.49 | 0.684 | 2.89E-04 | 0.00105 |
| uncultured_Barnesiella | 1.62 | 0.59 | 0.00627 | 0.0155 |
| uncultured_bacterium | -0.531 | 0.208 | 0.0111 | 0.0247 |
| Not_Assigned | 0.386 | 0.154 | 0.0126 | 0.0271 |
| unidentified | -1.3 | 0.555 | 0.0198 | 0.0408 |
| Aug21/Mar20 | | | | |
| p__Actinobacteriota | -1.74 | 0.285 | 2.03E-09 | 2.46E-08 |
| p__Verrucomicrobiota | -1.93 | 0.42 | 5.44E-06 | 3.11E-05 |
| p__Bacteroidota | -0.171 | 0.0733 | 0.0202 | 0.0479 |
| c__Coriobacteriia | -1.74 | 0.285 | 2.03E-09 | 2.90E-08 |

| | | | | |
|---|---|---|---|---|
| c__Verrucomicrobiae | -1.93 | 0.42 | 5.44E-06 | 3.39E-05 |
| c__Gammaproteobacteria | -1.15 | 0.294 | 1.13E-04 | 4.95E-04 |
| c__Bacilli | -1 | 0.364 | 0.00624 | 0.0177 |
| c__Bacteroidia | -0.171 | 0.0733 | 0.0202 | 0.0487 |
| o__Erysipelotrichales | -2.27 | 0.318 | 2.99E-12 | 6.22E-11 |
| o__Lactobacillales | -2.13 | 0.32 | 7.18E-11 | 1.38E-09 |
| o__Coriobacteriales | -1.74 | 0.285 | 2.03E-09 | 2.88E-08 |
| o__Peptococcales | -1.4 | 0.24 | 8.70E-09 | 1.07E-07 |
| o__Verrucomicrobiales | -1.93 | 0.42 | 5.44E-06 | 3.86E-05 |
| o__Burkholderiales | -1.15 | 0.294 | 1.13E-04 | 5.54E-04 |
| o__Acholeplasmatales | 1.24 | 0.513 | 0.0161 | 0.0396 |
| o__Bacteroidales | -0.171 | 0.0733 | 0.0202 | 0.0473 |
| f__Streptococcaceae | -2.61 | 0.221 | 6.61E-29 | 2.91E-27 |
| f__Erysipelotrichaceae | -2.27 | 0.318 | 2.99E-12 | 5.15E-11 |
| f__Atopobiaceae | -1.74 | 0.285 | 2.03E-09 | 2.87E-08 |
| f__Peptococcaceae | -1.4 | 0.24 | 8.70E-09 | 1.08E-07 |
| f__Akkermansiaceae | -1.93 | 0.42 | 5.44E-06 | 3.78E-05 |
| f__Lactobacillaceae | -1.49 | 0.325 | 5.74E-06 | 3.86E-05 |
| f__Sutterellaceae | -1.15 | 0.294 | 1.13E-04 | 5.66E-04 |
| f__Acholeplasmataceae | 1.24 | 0.513 | 0.0161 | 0.0405 |
| g__Lactococcus | -2.61 | 0.221 | 6.61E-29 | 3.66E-27 |
| g__Dubosiella | -2.31 | 0.215 | 1.56E-24 | 6.25E-23 |
| g__Coriobacteriaceae_UCG_002 | -1.74 | 0.285 | 2.03E-09 | 3.11E-08 |
| g__Ileibacterium | -1.94 | 0.328 | 6.13E-09 | 8.33E-08 |
| g__Faecalibaculum | -1.68 | 0.307 | 6.06E-08 | 6.92E-07 |
| g__Akkermansia | -1.93 | 0.42 | 5.44E-06 | 3.76E-05 |
| g__Lactobacillus | -1.49 | 0.325 | 5.74E-06 | 3.90E-05 |
| g__Blautia | -1.51 | 0.347 | 1.58E-05 | 9.63E-05 |
| g__Parasutterella | -1.15 | 0.294 | 1.13E-04 | 5.53E-04 |
| g__Bilophila | -0.955 | 0.271 | 4.64E-04 | 0.00184 |

| | | | | |
|---|---|---|---|---|
| g__Lachnoclostridium | -0.833 | 0.28 | 0.00306 | 0.00984 |
| g__Intestinimonas | -0.936 | 0.321 | 0.00369 | 0.0116 |
| g__A2 | -0.925 | 0.345 | 0.00764 | 0.0208 |
| g__Desulfovibrio | -0.734 | 0.281 | 0.00911 | 0.024 |
| g__Anaerotruncus | -0.813 | 0.333 | 0.0151 | 0.0372 |
| g__Anaeroplasma | 1.24 | 0.513 | 0.0161 | 0.0389 |
| g__Eubacterium_xylanophilum_group | -0.629 | 0.265 | 0.0181 | 0.0432 |
| g__Lachnospiraceae_UCG_006 | -0.709 | 0.301 | 0.0189 | 0.0451 |
| Lactococcus_lactis | -2.61 | 0.221 | 6.61E-29 | 3.27E-27 |
| Dubosiella_newyorkensis | -2.31 | 0.215 | 1.56E-24 | 6.19E-23 |
| Ileibacterium_valens | -1.94 | 0.328 | 6.13E-09 | 6.07E-08 |
| Lachnospiraceae_bacterium | -1.51 | 0.347 | 1.58E-05 | 7.62E-05 |
| Not_Assigned | 0.291 | 0.0739 | 9.16E-05 | 3.51E-04 |
| mouse_gut | -1.12 | 0.284 | 9.62E-05 | 3.59E-04 |
| Clostridium_leptum | 0.798 | 0.29 | 0.0061 | 0.0157 |
| uncultured_bacterium | -0.247 | 0.0999 | 0.0138 | 0.0307 |
| Aug21/May21 | | | | |
| p__Desulfobacterota | -1.27 | 0.294 | 1.94E-05 | 1.02E-04 |
| p__Proteobacteria | -1.11 | 0.39 | 0.00472 | 0.0129 |
| p__Deferribacterota | -1.32 | 0.537 | 0.0145 | 0.0365 |
| c__Desulfovibrionia | -1.27 | 0.294 | 1.94E-05 | 1.12E-04 |
| c__Alphaproteobacteria | -1.72 | 0.529 | 0.00119 | 0.00409 |
| c__Deferribacteres | -1.32 | 0.537 | 0.0145 | 0.0378 |
| o__Desulfovibrionales | -1.27 | 0.294 | 1.94E-05 | 1.38E-04 |
| o__Erysipelotrichales | -1.68 | 0.417 | 6.75E-05 | 4.05E-04 |
| o__Rhodospirillales | -1.72 | 0.529 | 0.00119 | 0.00452 |
| o__Oscillospirales | -0.87 | 0.322 | 0.00708 | 0.0215 |
| o__Deferribacterales | -1.32 | 0.537 | 0.0145 | 0.0386 |
| f__Streptococcaceae | -1.33 | 0.29 | 5.09E-06 | 4.03E-05 |
| f__Desulfovibrionaceae | -1.27 | 0.294 | 1.94E-05 | 1.37E-04 |

| | | | | |
|---|---|---|---|---|
| f__Erysipelotrichaceae | -1.68 | 0.417 | 6.75E-05 | 4.11E-04 |
| f__Tannerellaceae | -1.09 | 0.319 | 6.58E-04 | 0.00272 |
| f__uncultured | -1.72 | 0.529 | 0.00119 | 0.00444 |
| f__Bacteroidaceae | -0.699 | 0.233 | 0.00283 | 0.0092 |
| f__Ruminococcaceae | -0.991 | 0.352 | 0.00512 | 0.0155 |
| f__Oscillospiraceae | -0.905 | 0.324 | 0.00534 | 0.016 |
| f__Deferribacteraceae | -1.32 | 0.537 | 0.0145 | 0.038 |
| g__Bilophila | -1.89 | 0.356 | 1.63E-07 | 1.78E-06 |
| g__Lactococcus | -1.33 | 0.29 | 5.09E-06 | 3.67E-05 |
| g__Colidextribacter | -1.51 | 0.37 | 4.90E-05 | 2.80E-04 |
| g__Dorea | -1.22 | 0.341 | 3.99E-04 | 0.00167 |
| Not_Assigned | -0.829 | 0.249 | 9.27E-04 | 0.00341 |
| g__Incertae_Sedis | -1.08 | 0.331 | 0.00115 | 0.00409 |
| g__Ileibacterium | -1.37 | 0.431 | 0.00158 | 0.00535 |
| g__uncultured | -0.915 | 0.297 | 0.00218 | 0.00708 |
| g__Bacteroides | -0.699 | 0.233 | 0.00283 | 0.009 |
| g__Mucispirillum | -1.32 | 0.537 | 0.0145 | 0.0372 |
| g__Blautia | -1.08 | 0.456 | 0.0177 | 0.0442 |
| Lactococcus_lactis | -1.33 | 0.29 | 5.09E-06 | 3.15E-05 |
| Ileibacterium_valens | -1.37 | 0.431 | 0.00158 | 0.00512 |
| Clostridium_leptum | -0.954 | 0.381 | 0.0125 | 0.0288 |
| Lachnospiraceae_bacterium | -1.08 | 0.456 | 0.0177 | 0.0402 |
| Aug21/Nov22 | | | | |
| p__Actinobacteriota | -5.25 | 0.542 | 1.35E-20 | 5.69E-19 |
| p__Bacteroidota | 1.3 | 0.139 | 2.77E-19 | 8.73E-18 |
| p__Firmicutes | -1.82 | 0.329 | 4.99E-08 | 2.99E-07 |
| p__Desulfobacterota | -2.3 | 0.425 | 9.02E-08 | 5.17E-07 |
| p__Proteobacteria | -1.45 | 0.565 | 0.0107 | 0.0237 |
| c__Coriobacteriia | -5.25 | 0.542 | 1.35E-20 | 7.32E-19 |
| c__Bacteroidia | 1.3 | 0.139 | 2.77E-19 | 1.12E-17 |

| | | | | |
|---|---|---|---|---|
| c__Bacilli | -5.19 | 0.692 | 2.53E-13 | 3.72E-12 |
| c__Desulfovibrionia | -2.3 | 0.425 | 9.02E-08 | 6.09E-07 |
| c__Clostridia | -1.67 | 0.395 | 2.88E-05 | 1.11E-04 |
| c__Gammaproteobacteria | -2.32 | 0.559 | 3.82E-05 | 1.44E-04 |
| o__Erysipelotrichales | -7.23 | 0.604 | 1.73E-29 | 9.33E-28 |
| o__Coriobacteriales | -5.25 | 0.542 | 1.35E-20 | 5.23E-19 |
| o__Bacteroidales | 1.3 | 0.139 | 2.77E-19 | 9.35E-18 |
| o__Lactobacillales | -4.46 | 0.608 | 8.33E-13 | 1.18E-11 |
| o__Desulfovibrionales | -2.3 | 0.425 | 9.02E-08 | 6.96E-07 |
| o__Oscillospirales | -2.32 | 0.466 | 8.65E-07 | 5.31E-06 |
| o__Lachnospirales | -2.19 | 0.446 | 1.21E-06 | 7.13E-06 |
| o__Burkholderiales | -2.32 | 0.559 | 3.82E-05 | 1.69E-04 |
| o__Acholeplasmatales | -3.17 | 0.975 | 0.00123 | 0.00357 |
| f__Erysipelotrichaceae | -7.23 | 0.604 | 1.73E-29 | 7.61E-28 |
| f__Streptococcaceae | -4.84 | 0.419 | 7.96E-28 | 2.87E-26 |
| f__Atopobiaceae | -5.25 | 0.542 | 1.35E-20 | 4.13E-19 |
| f__Ruminococcaceae | -3.09 | 0.51 | 2.47E-09 | 2.65E-08 |
| f__Desulfovibrionaceae | -2.3 | 0.425 | 9.02E-08 | 7.94E-07 |
| f__Lachnospiraceae | -2.19 | 0.446 | 1.21E-06 | 7.52E-06 |
| f__Oscillospiraceae | -2.08 | 0.468 | 1.09E-05 | 5.92E-05 |
| f__Lactobacillaceae | -2.71 | 0.618 | 1.42E-05 | 7.50E-05 |
| f__Sutterellaceae | -2.32 | 0.559 | 3.82E-05 | 1.82E-04 |
| f__Butyricicoccaceae | -1.89 | 0.468 | 5.91E-05 | 2.66E-04 |
| f__Eubacterium_coprostanoligenes_group | -1.62 | 0.484 | 8.58E-04 | 0.00278 |
| f__Bacteroidaceae | 1.1 | 0.337 | 0.00116 | 0.00362 |
| f__Acholeplasmataceae | -3.17 | 0.975 | 0.00123 | 0.00377 |
| f__Muribaculaceae | 0.795 | 0.321 | 0.0137 | 0.0324 |
| g__Ileibacterium | -9.85 | 0.623 | 1.04E-46 | 1.49E-44 |
| g__Lactococcus | -4.84 | 0.419 | 7.96E-28 | 3.18E-26 |

| | | | | |
|---|---|---|---|---|
| g__Coriobacteriaceae_UCG_002 | -5.25 | 0.542 | 1.35E-20 | 4.24E-19 |
| g__Blautia | -4.88 | 0.659 | 5.08E-13 | 8.92E-12 |
| g__Incertae_Sedis | -3.51 | 0.479 | 8.65E-13 | 1.48E-11 |
| g__Desulfovibrio | -3.64 | 0.533 | 2.32E-11 | 3.33E-10 |
| Not_Assigned | -2.45 | 0.36 | 2.76E-11 | 3.89E-10 |
| g__Anaerotruncus | -4.28 | 0.633 | 3.67E-11 | 5.09E-10 |
| g__Faecalibaculum | -3.73 | 0.583 | 3.23E-10 | 4.15E-09 |
| g__Dubosiella | -2.39 | 0.408 | 8.46E-09 | 9.51E-08 |
| g__Lachnoclostridium | -3.05 | 0.532 | 1.58E-08 | 1.67E-07 |
| g__Intestinimonas | -3.25 | 0.61 | 1.46E-07 | 1.21E-06 |
| g__Lachnospiraceae_FCS020_group | -3.24 | 0.616 | 2.17E-07 | 1.74E-06 |
| g__Colidextribacter | -2.61 | 0.535 | 1.41E-06 | 8.59E-06 |
| g__Oscillibacter | -2.45 | 0.545 | 8.65E-06 | 4.35E-05 |
| g__Lachnospiraceae_UCG_001 | -3.57 | 0.801 | 1.01E-05 | 4.99E-05 |
| g__ASF356 | -2.61 | 0.589 | 1.09E-05 | 5.33E-05 |
| g__Lactobacillus | -2.71 | 0.618 | 1.42E-05 | 6.72E-05 |
| g__GCA_900066575 | -2.51 | 0.575 | 1.49E-05 | 6.99E-05 |
| g__uncultured | -1.86 | 0.43 | 1.86E-05 | 8.49E-05 |
| g__Dorea | -2.12 | 0.494 | 2.03E-05 | 9.19E-05 |
| g__Parasutterella | -2.32 | 0.559 | 3.82E-05 | 1.60E-04 |
| g__UCG_009 | -1.89 | 0.468 | 5.91E-05 | 2.39E-04 |
| g__Lachnospiraceae_UCG_006 | -2.11 | 0.573 | 2.58E-04 | 8.97E-04 |
| g__Tuzzerella | -1.91 | 0.521 | 2.74E-04 | 9.40E-04 |
| g__Bilophila | -1.88 | 0.515 | 2.84E-04 | 9.66E-04 |
| g__Eubacterium_coprostanoligenes_group | -1.62 | 0.484 | 8.58E-04 | 0.0026 |
| g__Bacteroides | 1.1 | 0.337 | 0.00116 | 0.00335 |
| g__Anaeroplasma | -3.17 | 0.975 | 0.00123 | 0.0035 |
| g__Roseburia | -2.13 | 0.673 | 0.00167 | 0.00457 |
| g__A2 | -1.72 | 0.656 | 0.00892 | 0.0202 |

| | | | | |
|---|---|---|---|---|
| g__Muribaculaceae | 0.78 | 0.321 | 0.0156 | 0.0332 |
| g__Lachnospiraceae_NK4A136_group | -1.3 | 0.542 | 0.0167 | 0.0354 |
| Aug21/Oct20 | | | | |
| p__Firmicutes | 1 | 0.322 | 0.0019 | 0.00557 |
| c__Clostridia | 1.2 | 0.387 | 0.00205 | 0.00659 |
| o__Oscillospirales | 1.38 | 0.456 | 0.00262 | 0.00864 |
| o__Lachnospirales | 1.06 | 0.436 | 0.0154 | 0.0404 |
| f__Muribaculaceae | 0.916 | 0.315 | 0.00376 | 0.0124 |
| f__Oscillospiraceae | 1.23 | 0.458 | 0.00729 | 0.022 |
| f__Ruminococcaceae | 1.31 | 0.499 | 0.00916 | 0.0269 |
| f__Lachnospiraceae | 1.06 | 0.436 | 0.0154 | 0.0407 |
| g__uncultured | 1.42 | 0.421 | 7.72E-04 | 0.00314 |
| g__Muribaculaceae | 0.928 | 0.315 | 0.00332 | 0.0114 |
| uncultured_Bacteroidales | 1.18 | 0.325 | 2.93E-04 | 0.0011 |
| uncultured_bacterium | 0.584 | 0.186 | 0.00175 | 0.00578 |

**Legend:** d (domain), p (phylum), c (class), o (order), f (family), g (genus), standard error (St. Error), false-discovery rate adjusted p-value (FDR)

**Table S14.** The full, shared, and unique contribution of each experimental factor to the variations in the gut microbiota composition. FDR-adjusted p-values <0.02 were considered significant. Full contribution represents how much all the experimental factors in our study explain the variations observed in the gut microbiome. Shared contribution represents how much each factor explains the variation without correcting for the other variables. Unique contribution is how much each factor explains the variations observed, after correcting for all other variables.

| | **Factor** | **Element** | ***P* value** | **Adj.r.squared** | **Percentage of variance (%)** | **Adj. *P* value** |
|---|---|---|---|---|---|---|
| 1 | full | full | NA | 0.488 | 48.800 | NA |
| 1 | Diet | shared | 0.001 | 0.233 | 23.300 | 0.001 |
| 2 | Sequencing batch | shared | 0.001 | 0.2091 | 20.910 | 0.001 |
| 3 | Compartment | shared | 0.001 | 0.128 | 12.800 | 0.001 |
| 4 | Animal Facility | shared | 0.001 | 0.089 | 8.900 | 0.001 |
| 5 | Genotype | shared | 0.001 | 0.051 | 5.100 | 0.001 |
| 6 | Age | shared | 0.001 | 0.047 | 4.700 | 0.001 |
| 7 | Angiotensin II treatment | shared | 0.001 | 0.028 | 2.800 | 0.001 |
| 8 | Sex | shared | 0.001 | 0.014 | 1.400 | 0.001 |
| 1 | Compartment | unique | 0.001 | 0.068 | 6.800 | 0.001 |
| 2 | Diet | unique | 0.001 | 0.060 | 6.000 | 0.001 |
| 3 | Sequencing batch | unique | 0.001 | 0.047 | 4.7 | 0.001 |
| 4 | Genotype | unique | 0.001 | 0.029 | 2.900 | 0.001 |
| 5 | Animal Facility | unique | 0.001 | 0.023 | 2.300 | 0.001 |
| 6 | Age | unique | 0.001 | 0.020 | 2.000 | 0.001 |
| 7 | Sex | unique | 0.001 | 0.004 | 0.400 | 0.001 |
| 8 | Angiotensin II treatment | unique | 0.001 | 0.004 | 0.400 | 0.001 |

**Legend**: Adj, adjusted; NA, not applicable.
